# ES-62 suppression of arthritis reflects epigenetic rewiring of synovial fibroblasts to a joint-protective phenotype

**DOI:** 10.1101/2020.10.08.331942

**Authors:** Marlene Corbet, Miguel A Pineda, Kun Yang, Anuradha Tarafdar, Sarah McGrath, Rinako Nakagawa, Felicity E Lumb, William Harnett, Margaret M Harnett

## Abstract

The parasitic worm product, ES-62 protects against collagen-induced arthritis, a mouse model of rheumatoid arthritis (RA) by suppressing the synovial fibroblast (SF) responses perpetuating inflammation and driving joint destruction. Such SF responses are shaped during disease progression by the inflammatory microenvironment of the joint that promotes remodelling of their epigenetic landscape, inducing an “aggressive” pathogenic SF phenotype. Critically, exposure to ES-62 *in vivo* induces a stably imprinted “safe” phenotype that exhibits responses more typical of healthy SFs. Surprisingly however, DNA methylome analysis reveals that rather than simply preventing the pathogenic rewiring of SFs, ES-62 induces further epigenetic remodelling, including targeting genes associated with ciliogenesis and differentiation, to program a distinct “protective” phenotype. Such unique behaviour signposts potential DNA methylation signatures predictive of pathogenesis and its resolution and hence, candidate mechanisms by which novel therapeutic interventions could prevent SFs from perpetuating joint inflammation and destruction in RA.

## Introduction

The Hygiene Hypothesis proposes that the recent, rapid eradication of parasitic worms and other pathogens occurring during urbanisation and industrialisation has resulted in populations exhibiting chronically unbalanced, hyperactive immune systems that likely contribute to the corresponding dramatically increased incidence of allergic, autoimmune and metabolic conditions worldwide^1^. Epidemiological evidence supports this hypothesis, most strikingly for autoimmunity in reports from India demonstrating clear evidence of an inverse relationship between infection with filarial nematodes and incidence of rheumatoid arthritis (RA) and systemic lupus erythematosus (SLE)^2–4^. Together with data from a range of experimental animal models of chronic inflammatory diseases^1^, these findings underpin the recent interest in developing “helminth therapies” based on infections with worms or their immunomodulatory products that have evolved to promote their survival by acting to dampen host inflammation and pathology^1^. Reflecting this, ES-62, a glycoprotein we discovered in the secretions of the filarial nematode *Acanthocheilonema viteae* that is immunomodulatory by virtue of multiple covalently attached phosphorylcholine moieties^1, 5^, can prevent disease initiation and progression in mouse models of asthma, dermatitis, RA, SLE and comorbidities arising from obesity-accelerated ageing^1, 6^.

RA is a chronic autoimmune inflammatory disease that targets articular joints, transforming the synovium into an inflamed, hyperplastic and invasive pannus that results ultimately in cartilage and bone destruction^7, 8^. Such joint destruction in RA and in mouse models of inflammatory arthritis (e.g. collagen-induced arthritis [CIA]), has long been associated with the chronic autoimmune inflammation that results from dysregulated T helper cell responses. Nevertheless, more recently, interest has focused on the role(s) that synovial fibroblasts (SFs) play in pathogenesis, from the early onset of the disease through to established inflammation and joint destruction and even the spread of disease to unaffected joints^7, 8^. During pathogenesis, SFs undergo an epithelial-mesenchymal transition (EMT)-like mechanism, adopting a mesenchymal/fibrotic phenotype^9–11^ that is epigenetically rewired to hyper-produce pro-inflammatory cytokines and drive joint destruction by secreting matrix metalloproteinases (MMPs) that damage cartilage and bone, and via RANKL, impact on osteoclastogenesis^7, 8, 12–15^. ES-62 prevents CIA by subverting TLR4 signalling to retune chronically elevated MyD88 levels to steady state, thereby resetting homeostatic immunoregulation and promoting resolution of auto-immune IL-17 (CD4^+^ and *γδ* T cell) responses primarily by restoring levels of regulatory B cells (Bregs)^16–19^. However, ES-62 also acts to suppress the aggressive hyper-inflammatory phenotype of SFs and prevent osteoclastogenesis^18, 20^.

In this study, our aim was to understand how SFs become rewired to the aggressive phenotype that drives joint pathology in the CIA mouse model and how ES-62 subverts this. We have therefore investigated the impact of the local pro-inflammatory environment pertaining during disease, specifically focusing on the signalling and epigenetic mechanisms by which IL-1*β* and IL-17 regulate the pathogenic phenotype of SFs. Reflecting their high levels of expression in the arthritic joint, we now show that chronic exposure to IL-1*β* and IL-17 *in vitro* can recapitulate such reprogramming, specifically the global DNA hypomethylation resulting from downregulation of DNA methyl-transferase 1 (DNMT1), and that both acute (cytokine and MMP production) and chronic (remodelling to a stable aggressive phenotype) SF responses to these pathogenic cytokines are dependent on ERK and STAT3 signalling. Our data reveal that ES-62 acts to rewire SFs away from their aggressive pathogenic phenotype during CIA, and this is associated with suppression of ERK and STAT3 activation and promotion of SOCS1/3 signalling. Critically however, ES-62 does not simply maintain/restore naïve cell status but induces additional methylome changes. Furthermore, this unique finding provides a first step towards identifying potential DNA methylation signatures predictive of pathogenesis and its resolution and hence, candidate mechanisms by which novel therapeutic interventions could prevent SFs from perpetuating inflammation and bone destruction in the joint.

## Results

We have previously shown that SFs from ES-62-treated CIA mice exhibit reduced spontaneous and IL-17-stimulated IL-6 release relative to those from PBS-treated CIA mice^18^. To investigate whether the ability of ES-62 to suppress the aggressive responses of SFs in CIA reflects stable rewiring of their functional phenotype, SF explant cultures (passaged for 3-4 weeks) from non-arthritic (Naïve; no CIA induction) and PBS- (CIA) and ES-62- (ES-62) treated mice undergoing CIA were examined for release of proinflammatory cytokines, chemokines and MMPs (Fig. 1). Such factors promote inflammation and degrade extracellular matrix, facilitating the hypoxia and loss of barrier integrity that drives cellular infiltration, pannus formation and joint destruction^21^. We now show that both the increased spontaneous release of IL-6 in CIA- relative to Naïve-SFs and the ES-62 rescue of CIA-SF responses back towards the naïve functional phenotype is evident even after such sustained culture of SFs, *ex vivo* (Fig. 1a). Moreover, IL-17-, IL-1*β*-, LPS- and BLP-stimulated release of IL-6 by SFs from PBS-treated CIA mice was similarly suppressed by *in vivo* ES-62 treatment (Fig. 1b-e), as was the spontaneous and IL-17- and LPS-stimulated CCL2 release by SFs from CIA mice (Fig. 1f & g).

**Fig. 1:**
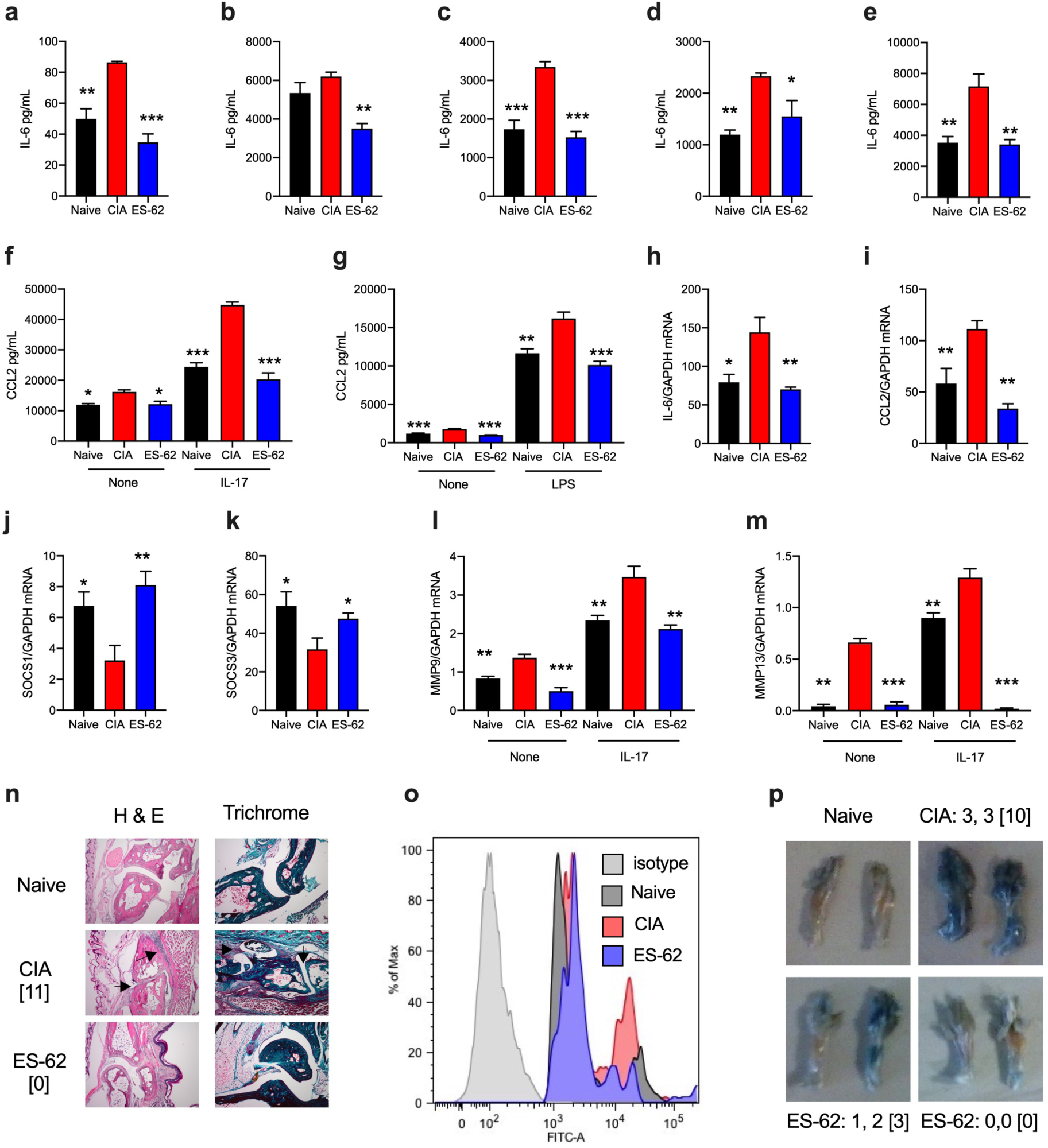
CIA induces an aggressive pathogenic SF phenotype that is counteracted by ES-62. 3-4 week explant SFs from naïve, CIA and ES-62-CIA (ES-62) mice were incubated overnight with (**a**) medium, (**b**) IL-17, (**c**) IL-1*β*, (**d**) LPS or (**e**) BLP and IL-6 release measured. Data are presented as mean (of means of triplicates) values ± SEM of n=3 independent cultures from single experiments representative of 3-7. Pooled data revealed that CIA-SFs showed 1.486 ± 0.180-fold (n=7, p<0.05) and ES-62-SFs, 0.873 ± 0.099-fold (n=4) spontaneous IL-6 release relative to Naïve SFs and IL-17-stimulated CIA-SFs exhibited x2.293 ± 0.566-fold (n=3, p<0.05) release compared to Naïve controls, whilst treatment with ES-62 reduced this hyper-production (x 1.871 ± 0.618-fold, n=3). (**f, g**) Spontaneous (None), IL-17- and LPS-stimulated CCL2 responses were elevated in CIA, relative to Naïve SFs and reduced in ES-62-SFs. Data are from single experiments representative of three. SFs were analysed for mRNA levels of (**h**) IL-6, (**i**) CCL2, (**j**) SOCS1 and (**k**) SOCS3, or following incubation overnight in medium (None) or IL-17, MMP9 (**l**) and MMP13 (**m**). Data are from single experiments representative of at least two and presented as mean (of means of triplicates) values ± SEM of n=3 independent cultures, assayed by qRT-PCR. (**n**) Joint pathology showing representative sections from mice, with total articular scores of the mouse examined [indicated]. Arrows show the cartilage erosion and cellular infiltration in CIA sections. (**o**) Flow cytometric analysis of the hypoxic status of joint SFs determined by pimonidazole *in vivo*. (**p)** Representative images of joint vascular leakage (Evan’s Blue), where articular score for individual paws and [total mouse score] are from one experiment representative of two independent models. Throughout, SFs were pooled from individual mice to generate representative explant cohorts with articular scores: (**a-e, g, k**) CIA, 3.17 ± 1.38, n=6; ES-62, 0.5 ± 0.22, n=6; (**f**) CIA, 8.5 ± 1.19, n=4; ES-62, 0.8 ± 0.49, n=5; (**h-j**, **l, m)** CIA, 3.66 ± 1.5, n=6; ES-62, 1.5 ± 1.15, n=6; (**o**) CIA, 4.667 ±1.99, n=6; ES-62, 0.8 ± 0.8, n=5. In all panels, *p<0.05; **p<0.01; ***p<0.001 relative to CIA SFs.

The pattern of release of inflammatory cytokines and chemokines was mirrored at the mRNA level with ES-62 reducing the elevated expression of IL-6 and CCL2 exhibited by CIA-SFs again back to those observed in SFs from naïve mice (Fig. 1h & i). SOCS proteins are key negative regulators of proinflammatory cytokine signalling, cell proliferation and migration that have been reported to exhibit reduced expression in RA joint cells, including synoviocyte-like fibroblasts^22–25^. Thus, consistent with their aggressive inflammatory phenotype, SOCS1 and SOCS3 are downregulated in CIA-SFs, whilst exposure to ES-62 rescues their expression (Fig. 1j & k). Moreover, in addition to their hyper-production of cytokines and consistent with their role in joint destruction, CIA-SFs also generate elevated levels of MMP9 and MMP13 in a spontaneous and IL-17-stimulated manner and again, treatment with ES-62 acts to suppress these (particularly MMP13) pathogenic effectors (Fig. 1l & m). Reflecting the increased pro-inflammatory and cartilage/extracellular matrix-degrading SF activities, the joint damage observed in CIA-mice is associated with increased levels of hypoxia in joint cells and also induction of vascular leakage: however, in each case this pathology is countered by ES-62 (Fig. 1n-p).

Collectively, these data suggested that prophylactic treatment with ES-62 prevented/ reversed the transformation of SFs to an aggressive pathogenic phenotype. Recent studies have revealed that this transformation is associated with epigenetic rewiring of the SFs, with much interest focusing on their global DNA methylation status due to the association of promoter region hypo-methylation with the de-repression of inflammatory genes driving SF transformation^7, 8, 12–15^. We therefore investigated whether naïve and CIA-SFs exhibited differential global DNA methylation and whether any changes could be mimicked by sustained *in vitro* treatment of naïve SFs with IL-17 or IL-1*β*, cytokines that synergistically drive joint inflammation and damage^26, 27^. As predicted, CIA-SFs exhibited hypo-methylated global DNA relative to those from naïve mice and this could indeed be mirrored by sustained *in vitro* treatment (for 2 weeks) of naïve SFs with either IL-17 or IL-1*β* (Fig. 2a). The functional relevance of this chronic cytokine-induced hypomethylation is illustrated by the associated spontaneous and stimulated (IL-17-, IL-1*β*- and LPS/TLR4-) hyper-elevated IL-6 responses, particularly following chronic treatment with IL-1*β* (Fig. 2b-d), that are reminiscent of those seen in CIA- relative to naïve SFs. Global DNA hypomethylation is associated with reduced expression of DNA methyltranferase-1, (DNMT1) and consistent with this, we found reduced expression of this enzyme in CIA-SFs (Fig. 2e & f) and IL-1*β*- and, to a lesser extent, IL-17-treated SFs (Fig. 2g), compared to their control naïve cohorts. Moreover, exposure to the DNMT1 inhibitor, 5-azacytidine (5-aza; 1 week) reduces global DNA methylation in naïve SFs and again, this is associated with increased spontaneous and stimulated IL-6 secretion and MMP13 expression (Fig. 2h-j), providing corroborative evidence for this mechanism of cytokine-induced SF pathogenesis in CIA.

**Fig. 2.**
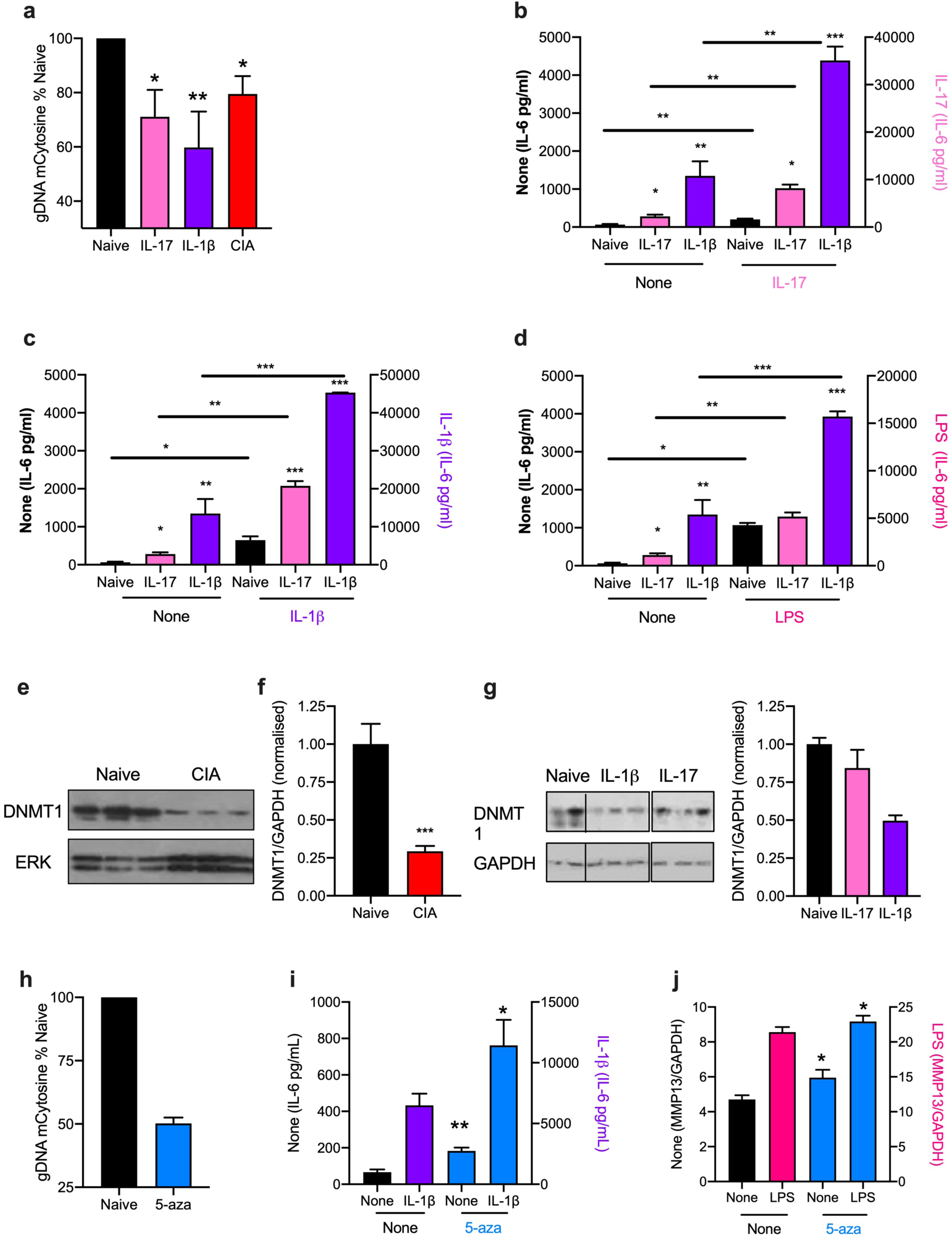
IL-1β and IL-17 rewire naïve SFs to a CIA-like phenotype. (a) analysis of global methylation status of naïve SFs incubated in medium alone (Naïve) or chronically treated with IL-17 or IL-1*β*, compared to that of CIA-SFs. Data are pooled from 3-5 independent experiments and are means ± SEM presented as percentage of the Naïve sample, where *p<0.05 and **p<0.01 relative to Naïve SFs. The CIA SFs analysed in these independent experiments were explanted from 3 independent models where the mice exhibited articular scores of 3.6 ± 1.5, n=6; 4.5 ± 1.3, n=10 and 2.5 ± 0.8, n=12. (**b-d)** Naïve SFs incubated in medium alone (Naïve) or chronically treated with IL-17 or IL-1*β* were re-challenged with medium (None; **b-d**), or medium containing IL-17 (**b**), IL-1*β* (**c**) or LPS (**d**) and levels of IL-6 release, determined and presented in separate panels for clarity. Data are presented as means ± SEM of n=3 independent cultures, where *p<0.05, **p<0.01 or ***p<0.001 are relative to the relevant Naïve SF control group (same data **b-d**). Data are from a single experiment although those in **b** are representative of 2 independent experiments. (**e-f**) Western blot analysis (images and quantitation) of DNMT1 expression in Naïve and CIA (articular score 3.6 ± 1.5, n=6) SFs (**e**: each lane represents independent cultures; **f**: quantitation using 2 independent experiments where ***p<0.001 for expression in CIA- versus naïve-SFs, both n=7 cultures). (**g)** Western blot analysis (images and combined quantitation) of DNMT1 expression in naïve SFs incubated in medium alone (Naïve) or chronically treated with IL-17 or IL-1*β* where the lanes represent individual replicate cultures from a single experiment. **(h-j)** Naïve SFs were chronically incubated in medium alone (None) or containing 5- aza and then assessed for their **(h)** global DNA methylation status, **(i)** IL-6 release in response to IL-1*β* challenge and **(j)** MMP13 mRNA levels in response to LPS-stimulation. Data are presented as means ± SEM, where n=3 independent cultures (**h** and **i**) or means ± SD, n=3 triplicate analyses (**j**) and *p<0.05 or **p<0.01 relative to the appropriate stimulus control (“None”).

To address identifying the upstream signalling mechanisms involved in driving such global DNA hypomethylation in CIA-SFs, we focused on the role of the ERK MAPkinase and STAT3 pathways, already implicated in IL-17- and IL-1*β−* (directly and indirectly via IL-6 signalling) driven SF responses and RA pathogenesis^28–33^. Here we confirm by a combination of flow cytometric, FACE and Western blot assays that IL-17 acutely signals via ERK and STAT3 (increasing the pERK/ERK and pSTAT3/STAT3 expression ratios) in naïve SFs (Fig. 3a-d). These studies also demonstrated that whilst CIA-SFs appear to exhibit stronger and/or more prolonged ERK and STAT3 activation than naïve SFs in response to IL-17, the low steady-state levels of ERK and STAT3 activation are comparable in SFs from naïve and CIA mice (Fig. 3c & d). Moreover, exposure of CIA-mice to ES-62 suppresses this ERK and STAT3 (the latter back towards basal levels at 10 mins post-stimulation) hyper-responsiveness (Fig. 3c & d), suggesting that in addition to reducing IL-17 levels *per se*, ES-62 can also block at least some of its subsequent downstream actions. Sustained activation of ERK and STAT3 signalling (Fig. 3e) was also detected following chronic stimulation with IL-17 or IL-1*β*, under the conditions associated with induction of global DNA hypomethylation and epigenetic rewiring (Fig. 2). However, reflecting the comparable steady-state levels observed in naïve and CIA-SFs (Fig. 3c & d), treatment of naïve SFs with 5-aza to induce DNA hypomethylation does not impact on basal ERK and STAT3 signalling (Fig. 3f): rather, these data suggest that SF rewiring licenses hyper-responsiveness of these signalling pathways to chronic inflammatory mediators. Consistent with roles for ERK and STAT3 signalling in SF pathogenesis, the STAT3 inhibitor 5.15DPP (iSTAT3; Fig. 3g) and MEK inhibitor PD98059 (iERK; Fig. 3h) not only suppress acute IL-17-stimulated IL-6 and MMP9 and MMP13 expression, but also prevent chronic IL-17- and IL-1*β*-mediated hypomethylation of global DNA (Fig. 4a-d). Further supporting a role for IL-1*β* in SF pathogenesis, co-treatment of CIA mice with IL-1*β* abolishes the ability of ES-62 to protect against development of joint inflammation and destruction: rather, these animals exhibit the exacerbated disease seen with CIA-mice treated with IL-1*β* alone (Fig. 4e-g) and that is associated with hyper-pathogenic SF MMP responses (Fig. 4h & i) that likely contribute to the enhanced joint damage observed in the IL-1*β*-treated CIA mice.

**Fig. 3.**
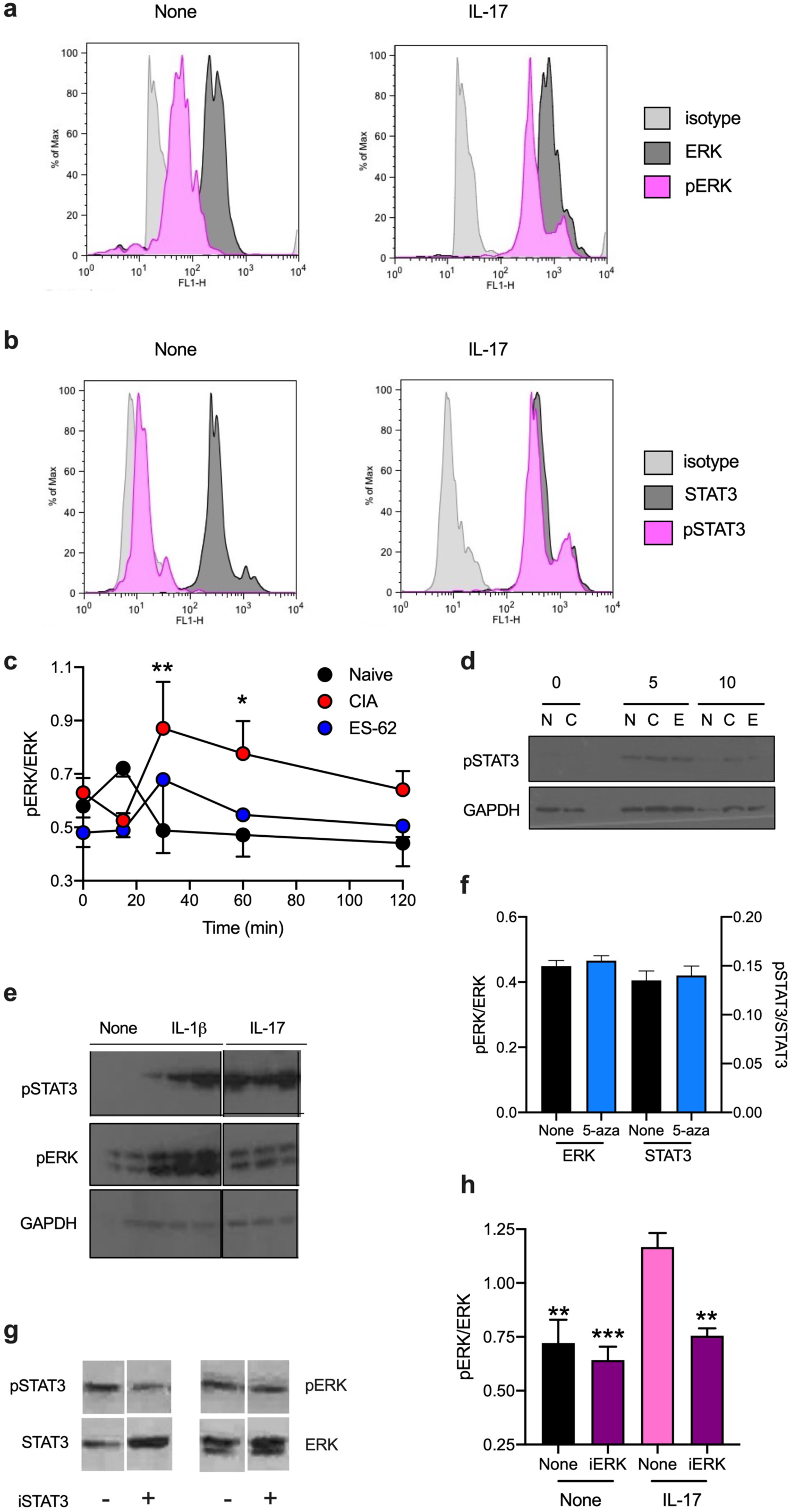
IL-17 and IL-1β stimulate ERK and STAT3 activation in SFs. Flow cytometric analysis of (**a**) ERK and (**b**) STAT3 activation, determining the relative expression of dually phosphorylated versus total ERK or phosphorylated STAT3 versus total STAT3 expression following incubation of Naive SFs in medium alone or containing IL-17 for 20 min. (**c**) SFs from Naïve, CIA- (articular score; 2.8 ± 1, n=7) and ES-62-CIA (articular score; 0.8 ± 0.8, n=5) mice were stimulated with IL-17 for the indicated times and ERK activation measured by the pERK/ERK ratio, determined by FACE assay: data are expressed as means ± SEM, n=3 independent cultures and are representative of two independent experiments, *p<0.05 and **p<0.01 for CIA versus Naïve SFs. (**d**) SFs from naïve (N), CIA- (C) and ES-62-CIA (E) mice were analysed for expression of pSTAT3 following stimulation with IL-17 for the indicated times (min) and for the data shown, each lane represents an individual culture. (**e**) Naïve SFs chronically incubated in medium alone (None) or containing IL-17 or IL-1*β* were analysed for pERK, pSTAT3 and GAPDH expression by Western blot: individual lanes represent independent SF cultures. (**f**) Naïve SFs chronically incubated in medium (None) or containing the DNMT1 inhibitor, 5-aza were subsequently analysed for their ERK and STAT3 activation status by FACE assay. Data are presented as the mean ± SEM, n=3 independent cultures. **(g)** Naïve SFs were pre-incubated with the STAT3 inhibitor 5.15 DPP (iSTAT3; 50 µM, +) or medium (-) for two hours prior to stimulation for 20 min with IL-17 and STAT3 and ERK activation assessed by Western blot analysis of pSTAT3, STAT3, pERK and ERK expression. (**h**) Preincubation with the MEK inhibitor, PD98059 (iERK; 25 µM) for 2 h blocked IL-17-stimulated ERK activation in Naïve SFs as determined by analysis of the pERK/ERK ratio measured by the FACE assay and where data represent mean values ± SEM of n=3 independent cultures and **p<0.01 and ***p<0.001, relative to the IL-17-stimulated control (“None”) sample.

**Fig. 4.**
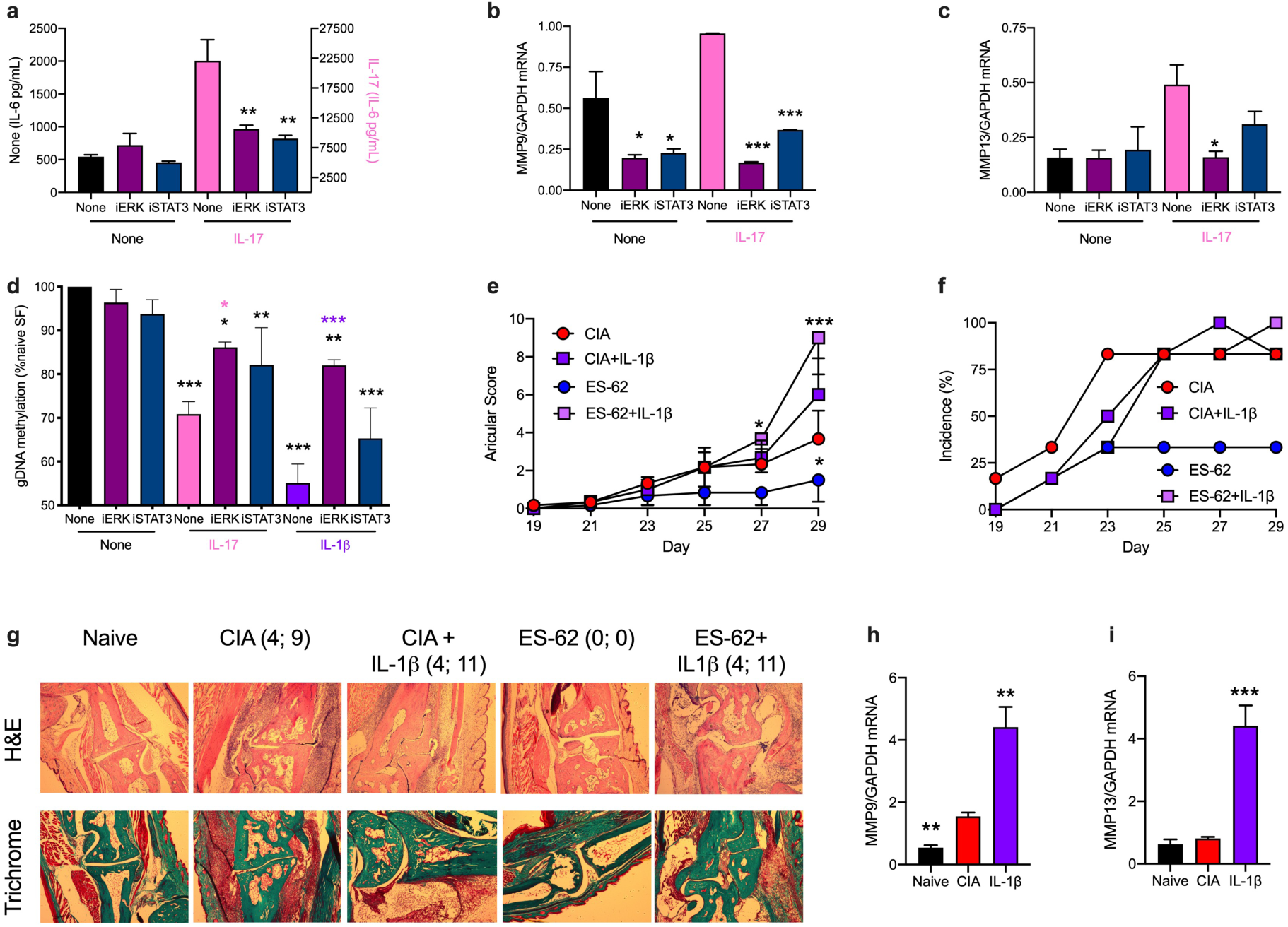
ERK and STAT3 signalling is required for IL-17 and IL-1β-driven generation of a CIA-like phenotype in Naïve SFs. (a-c) Naïve SFs were pretreated with iERK or iSTAT3 for two hours priors to incubation in medium alone (None) or with IL-17 overnight and levels of IL-6 release (**a**) or MMP9 and MMP13 mRNA (**b** & **c**) measured. (**d**) Naïve SFs pretreated with iERK and iSTAT3 for two hours were chronically incubated in medium alone (None) or containing IL-17 or IL-1*β* and then their global DNA methylation status assessed. Data are shown as mean values ± SEM, n=3 independent cultures and where *p<0.05, **p<0.01 and ***p<0.001 relative to appropriate None control (**a-c**) and None/None control (**d**) and where *p <0.05 is relative to its IL-17 control and ***p<0.001 is relative to its IL-1*β* control. CIA was induced in mice treated with PBS (CIA) or ES-62 on days -2, 0 and 21 and then with PBS or IL-1*β* (1 µg dose) twice-weekly from day 21 and articular scores (**e**) and incidence of pathology (**f**) monitored. Articular score data (**e**) are presented as mean values ± SEM with n=6 mice/group where *p<0.05 or ***p<0.001 are relative to the CIA group and incidence (**f**) represents the % mice displaying scores >1. Representative sections (**g**) of joint pathology from all groups are shown with the score of the joint displayed and total articular score of the mouse presented as indicated. MMP mRNA levels (**h, i;** articular scores, CIA: 3.66 ± 1.5, n=6; CIA + IL-1*β*: 6 ± 1.93, n=6) are presented as mean values ± SEM for n=3 independent cultures and **p<0.05 and ***p<0.001 are relative to the CIA group.

Non-coding RNAs, particularly microRNAs (miRs) have increasingly been implicated in bridging cytokine and TLR signalling (specifically JAK/STAT and MAPkinase pathways) with epigenetic rewiring and consequent transcriptional reprogramming: in turn, the dysregulated expression of certain miRs has been attributed to epigenetic changes present in chronic inflammatory disease^34–40^. Indeed, a wide range of miRs have been implicated in development of RA, with miR-146 and miR-155 proposed to be major species associated with SF pathogenesis, both being elevated in fibroblast-like synoviocytes from RA patients (RA SFs) relative to osteoarthritis (OA) controls and associated with disease activity/severity^35, 36, 38, 39^. We therefore screened a range of miRs implicated in regulating IL-1R/TLR and cytokine signalling and/or exhibiting modulated expression levels in CIA and human RA. MiRs-19b, -23b, -34a, -146 and -155 were found to be expressed in SFs from naïve and CIA-mice and reflecting previous reports that they are elevated in RA SFs and by association implicated in their pathogenesis^41^, miR-146 and -155 levels were found to be higher in CIA than naïve SFs (Fig. 5a). Moreover, and perhaps consistent with the proposed pro-inflammatory roles of TLR signalling in CIA/RA pathogenesis^42, 43^, LPS/TLR4 signalling substantially increases miR-155 levels in naïve SFs and this increase is associated with similar kinetics of MyD88 and IL-6 (although not Traf6) upregulation, at the mRNA level (Fig. 5b). We therefore investigated whether the (TLR4/MyD88-dependent) protective effects of ES-62 against pathogenic TLR and cytokine signalling and consequent joint damage were associated with modulation of expression of our panel of SF-associated miRs in CIA-SFs back towards the levels found in naïve SFs: this revealed that ES-62 indeed acted to reduce the elevated levels of miR-19b, -146 and -155 pertaining in CIA SFs (Fig. 5c-g). Interestingly, however, although miR-155 has been associated with driving pro-inflammatory innate and adaptive immune responses in RA and the CIA mouse model of RA^40, 44, 45^, its role(s) in SF pathogenesis is unclear due to the contradictory nature of the data in the literature^35, 36, 38, 39, 46^. Thus, for example, early studies perhaps surprisingly reported that miR-155 plays a joint-protective role in RA SFs, having been associated with downregulation of MMP1 and MMP3, mediators that promote SF proliferation and invasion, as well as bone destruction^35, 36, 38, 39, 46^. However, although these studies also indicated that miR-155 did not impact on MMP9 or MMP13 expression or modulate spontaneous or stimulated (TLR ligands/cytokines) TGF*β* or IL-6 production^41, 47^, more recent work has identified a key role for this miR in driving (spontaneous and TNF*α*-stimulated) pathogenic responses (proliferation as well as IL-1*β* and IL-6 production) via downregulation of its target FOXO3^48–50^, which acts to suppress these functional outcomes in RA-SFs^51^. Thus, given the interactions of miR-155 with multiple TLR and cytokine (IL-1*β*, IL-10, IL-17, TNF*α*, GM-CSF) pathways implicated in regulating arthritogenesis^38, 39^, we further investigated the role of this element in SF pathogenesis and ES-62-mediated protection. Firstly, we investigated whether miR-155 drove pro-inflammatory cytokine production in our SF system by exploiting the miR-155-LacZ reporter strain of mice, in which replacement of the miR-155 coding sequence by a LacZ cassette generates a non-functional allele^52, 53^: naïve SFs from wild type and miR155-deficient (LacZ heterozygotes and homozygous null) mice showed that basal and IL-17-, IL-1*β*- and LPS-stimulated IL-6 and CCL2 release was substantially dependent on miR-155 expression (Fig. 5h & i). Consistent with this, enforced expression of miR-155 via transfection with miR-155-5p mimic, increased basal release of IL-6 and CCL2 by SFs from each of Naïve, CIA and CIA-ES-62 mice and this was associated with profound suppression of SOCS1 (Fig. 5j-l), as previously reported for the association between high levels of miR-155 and IL-1*β* and TNF*α* in peripheral blood from RA patients^54^. However, this acute treatment with exogenous miR-155 did not impact on the expression of DNMT1 in any of the cohorts of SFs (Fig. 5m), reflecting the finding that its enforced expression did not convert either Naïve or ES-62 SFs to the hyper-responsive CIA phenotype, at least in terms of IL-6 and CCL2 release (Fig. 5j & k), but rather appeared to globally enhance cytokine release, perhaps in part by acutely suppressing SOCS1 that can target TLR signalling^22^. Surprisingly, however, these studies additionally showed that rather than preventing DNMT1 downregulation and consequently, global DNA hypomethylation to maintain the Naïve SF phenotype, ES-62 appeared to further reduce its expression suggesting that instead of preventing epigenetic rewiring, it enhanced it. Again, this was not impacted by enforced expression of miR-155 (Fig. 5m).

**Fig. 5.**
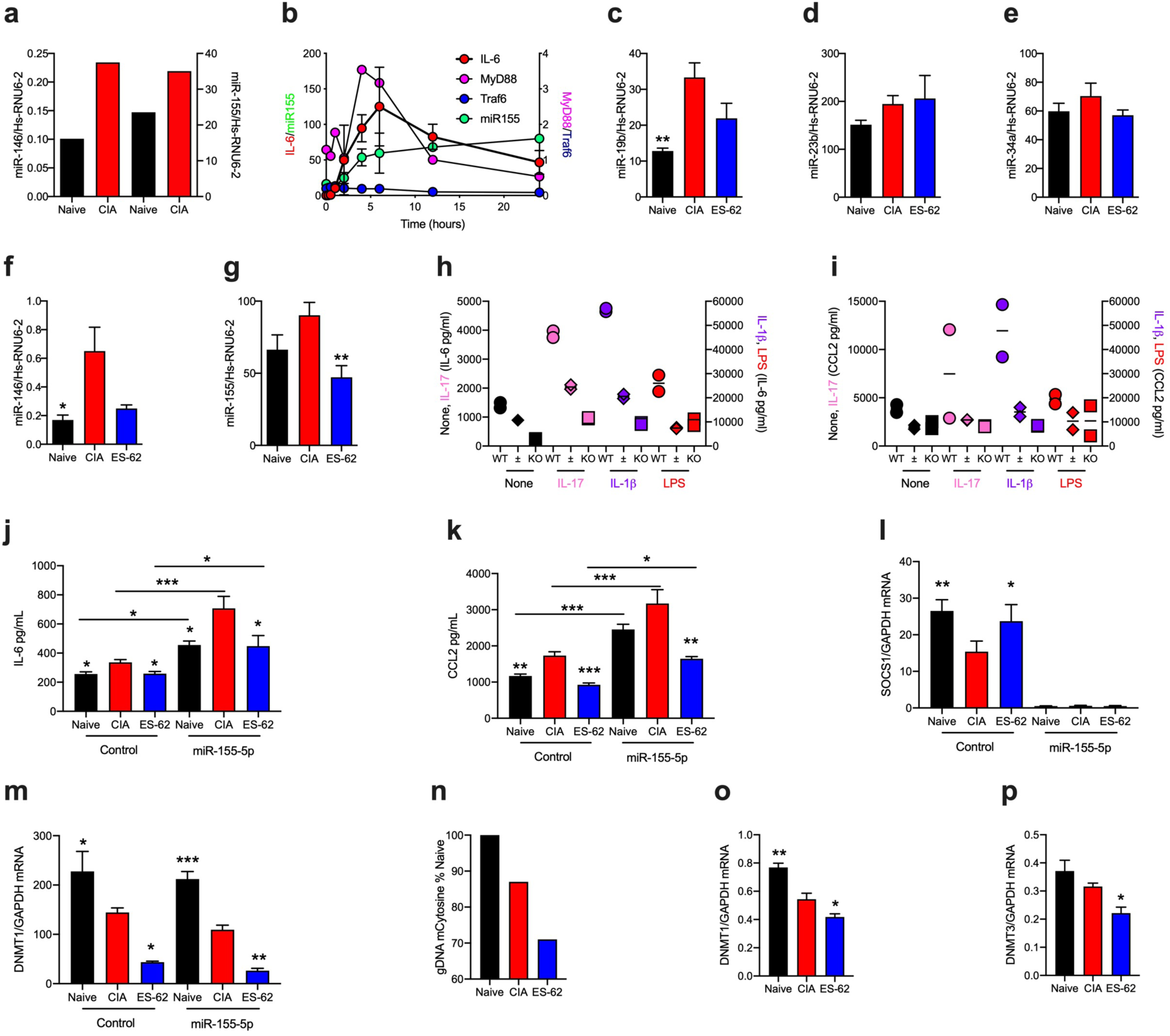
miR-146 and -155 are potential targets of ES-62 in suppressing the aggressive CIA-SF phenotype. (a) SFs from Naive and CIA-mice were assessed for their expression of miR-146 and miR-155 and data are presented from a single experiment. (**b**) Naïve SFs were stimulated with LPS for the indicated time and levels of miR-155, MyD88, Traf6 and IL-6 mRNA levels measured. Data are presented as mean values ± SEM, n=3 experiments for IL-6 and single experiments for the other mediators. (**c-g**) Expression levels of the indicated miRs were determined for SFs from Naïve, CIA (articular score: 4.67 ± 2, n=6) and ES-62-CIA (0.8 ± 0.8, n=6) mice stimulated with LPS for 6 h and data presented as mean values ± SEM, n=4-6 independent cultures and where *p<0.05 and **p< 0.01 relative to the CIA group. Although miR-19b or- 146 expression in ES-62-CIA SFs was not significantly reduced (p=0.06 and 0.07, respectively) when compared to that of CIA-SF, it was not significantly elevated when compared to that in the naïve SF controls. **(h & i)** SFs from wild type and miR-155 heterozygous (±) and homozygous (KO) C57BL/6 mice were incubated with the indicated stimuli and IL-6 **(h)** and CCL2 **(i)** release measured. Data shown represent the mean values (of ELISA triplicates) for two mice of each phenotype. **(j-m)** SFs from Naïve, CIA- (articular score 3.17 ± 1.38, n=6) and ES-62-CIA- (articular score 0.5 ± 0.22, n=6) mice were transfected for 24 hours with miR-155-5p mimic (50 nM) or control miR and then IL-6 and CCL2 release **(j & k)** and SOCS1 and DNMT1 mRNA levels **(l & m)** assessed. Data are from a single experiment and are expressed as the mean values (of triplicate assays) ± SEM, n= three independent cultures and where *p<0.05; **p<0.01, *** p<0.001. The IL-6 and CCL2 data were validated in an independent set of three further cultures. SFs from naïve, CIA (articular score 3.66 ± 1.5, n=6) and ES-62 CIA (articular score 1.5 ± 1.15 n=6) mice were assessed for their levels of (**n**) global DNA methylation, (**o & p)** mRNA levels of DNMT1 and DNMT3 expression, where data are shown as mean values ± SEM, n=3 independent cultures where *p<0.05 and **p<0.05 are relative to Naïve SFs.

We therefore next determined whether ES-62 could also induce further global DNA hypomethylation in CIA SFs, rather than block the epigenetic remodelling observed during pathogenesis, and this indeed proved to be the case (Fig. 5n). Moreover, although DNMT1 has generally been thought to be responsible for maintenance of DNA methylation during cell division, it is increasingly evident that rather than being restricted to *de novo* imprinting roles during development, DNMT3 is an important contributor to the maintenance of the DNA methylome landscape^55^. Of relevance therefore, parallel mRNA expression analysis showed that whilst rewiring of SFs from CIA mice was reflected by reduced levels of DNMT1 but not DNMT3, the more pronounced remodelling of the methylome seen with CIA-ES-62 SFs was associated with downregulation of both DNMT1 and DNMT3 (Fig. 5o & p).

Collectively, these data suggested that rather than blocking pathogenic transformation of SFs, ES-62 further epigenetically modified these to a “resolving” phenotype. To address determining whether this is the case, we undertook methylome analysis of naïve, CIA and ES-62-CIA SFs (Fig. 6) and indeed heatmap analysis of the methylation percentage of the Promoter 2K region (2000 bp upstream of TSS) appears to support this hypothesis as hierarchical clustering revealed the profile of ES-62-CIA SFs to be distinct to those of both the (more closely related) naïve and CIA groups (Fig. 6a). Moreover, deeper analysis of the differential (top 100 sites) methylation of Promoter 3K (3000 bp upstream of transcription start site [TSS]), TSS (1000 bp upstream to 1000 bp downstream of TSS) and gene body (start of 5’UTR to end of 3’UTR) regions, comparing (1) CIA relative to naïve, (2) ES-62-CIA relative to naive and (3) ES-62-CIA relative to CIA, SFs, indicated that ES-62 only acted to restore the methylation status back towards that seen in SFs from naïve mice for a limited number of genes at each of the gene regions. Rather, it induced further methylation changes, differentially inducing both hypo- and hyper-methylation of various genes relative to their naïve and CIA counterparts in each gene region examined (Fig. 6b). The functional outcome of methylation status at these various gene sites is not straightforward to understand however, as whilst hypermethylation at the promoter, TSS and 1st Exon sites is associated with gene silencing, within the gene body it appears to correlate with gene expression and regulation of splice variants^56–59^. In support of this, visualization of the differential profiles of CpG methylation at exon 3 of the ES-62 target, MyD88 (Fig. 6c) shows hypermethylation, and hence potential upregulation, in CIA SFs relative to that in naïve and ES-62-CIA SFs. Such hypermethylation could also conceivably favour expression of the pro-inflammatory long form splice variant of MyD88 over the dominant negative, anti-inflammatory short form splice variant (exon 2 excision)^60–62^ during CIA. Moreover, and consistent with our functional data showing that whilst CIA reduces SOCS1 expression, exposure to ES-62 restores it to at least the levels found in naïve SFs, visualization of the differential profiles of CpG methylation within the Promoter and TSS regions of the SOCS1 gene shows that CIA-SFs display hypermethylation (gene silencing) relative to SFs from both Naïve and ES-62-CIA mice (Fig. 6d).

**Fig. 6.**
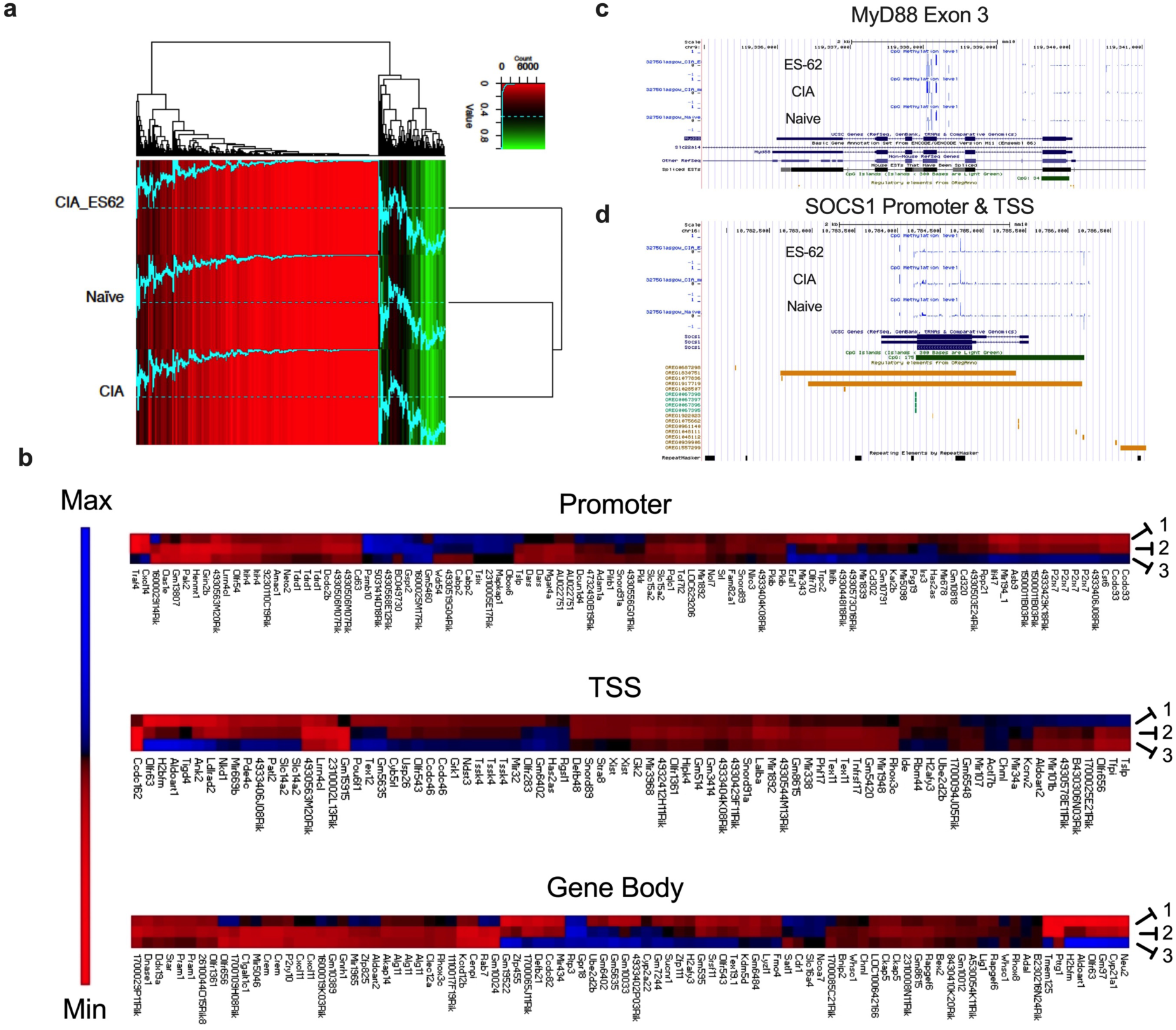
ES-62 induces a DNA methylation phenotype that is distinct from that of both Naïve and CIA-SFs. RRBS DNA methylome analysis was performed on SFs representative of the Naïve (n=6), CIA (articular score 3.17 ± 1.38, n=6) and ES-62-CIA (articular score 0.5 ± 0.22, n=6) cohorts, **(a)** A heatmap of methylation percentages of “Promoter2K” regions (0-2,000bp upstream of TSS) for all cohorts. Rows are hierarchically clustered using the Euclidean distance metric. Bright red indicates 0% methylation, black indicates 50% methylation, and bright green indicates 100% methylation. The blue line corresponds to the methylation percentage of the respective regions, thus facilitating visual comparison of methylation percentages across samples (the dotted line represents 50% methylation for reference). (**b**) Methylome analysis of SFs from Naive, CIA- and ES- 62-CIA-mice showing Heat Map analysis of the top 100 differentially methylated loci in the promoter/enhancer (Promoter 3K; left panel), transcription start site (TSS; middle panel) and core gene body (Core; right panel) regions of total genomic DNA. In each case, hypo (red) or hyper (blue) methylation in SFs from PBS- or ES-62- treated CIA-mice relative to SFs from healthy mice is shown in lanes 1 and 2 respectively, whilst that of SFs from ES-62- relative to PBS-treated CIA-mice is shown in lane 3. (**c & d**) Visualisation of the exon 3 region of MyD88 (**c**) and Promoter/TSS regions of SOCS1 (**d**) showing their differential CpG methylation in Naïve, CIA and ES-62-CIA cohorts with images created by uploading BED files in the UCSC browser.

As a first step to identifying key genes targeted by ES-62 in its differential epigenetic rewiring of pathogenic SFs in CIA we adopted a binary approach, examining sites that were either essentially fully methylated (>90%) or demethylated (0%) throughout the various gene regions for each of the treatment groups. Firstly, pathway analysis (using String software) of the fully methylated genes predicted quite distinct interactions/pathways (clusters) of functionally disparate classes of genes (summarised in pie-charts) for every treatment group in each of the Promoter 3K (Fig. 7a-c), TSS (Fig. 7d-f), 1^st^ Exon (start of 1^st^ exon to end of 1^st^ intron of gene; Fig. 7g-i) and Gene Body regions (Fig. 7j-l). For example, the “silenced” (hypermethylated) genes identified in the Promoter 3K and TSS regions of the naïve group (Fig. 7a & d) were predominantly epigenetic elements (chromatin and histone modifiers) and differentiation (development/oncofetal reprogramming factors) and ubiquitin (regulation of protein stability and trafficking) pathway components. By contrast, and reflecting their remodelled “transformed” phenotype, many of these elements were hypomethylated in SFs from the CIA group (Fig. 7b & e), which instead displayed hypermethylation of signalling, metabolic and neuropeptide genes (e.g. Fabp7, Mapk10, Mcr2 [ACTH-R], Gip [incretin], Cort) that are associated with suppression of inflammation, cell migration and promotion of bone formation. Moreover, exposure to ES-62 resulted in hypermethylation of genes associated with developmental reprogramming and ciliogenesis (Fig. 7c & f), cilia being sensory organelles which play roles in development and tissue repair/regeneration, coordinating cell migration, differentiation and signal transduction responses. Interestingly therefore, given that “aggressive” SFs display a hyperplasic, invasive phenotype, aberrant ciliogenesis and/or dysregulation of its functional outcomes is associated with cancer and chronic inflammatory and metabolic disorders^63–68^.

**Fig. 7.**
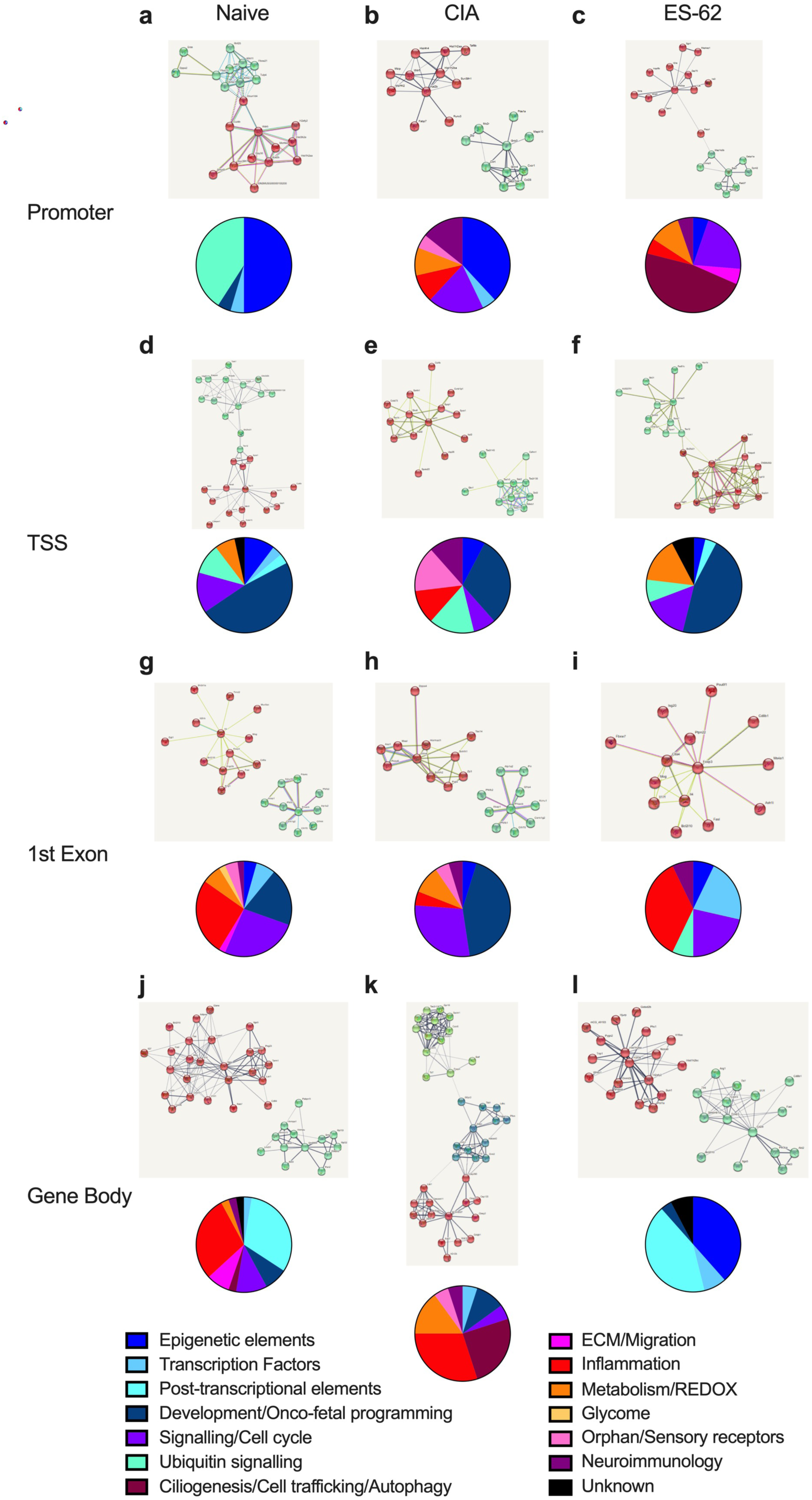
String pathway analysis of genes displaying fully methylated (>0.9) CpG regions at various sites. Genes displaying >0.9 methylation of CpG sites in the Promoter 3K (**a-c**), TSS (**d-f**), 1^st^ Exon (**g-i**) and Gene body (**j-l**) regions were modelled by String pathway analysis software: the clusters of differential pathway interactions predicted for the Naïve (**a, d, g, j**), CIA (**b, e, h, k**) and ES-62-CIA (ES-62; **c, f, i, l**) cohorts and the functional classification of the genes involved, summarized by pie-chart analysis, are shown.

Moreover, cilia have also been implicated in coordinating the crosstalk between the ubiquitin/proteosomal and autophagolysosomal systems and we have recently shown^69^ that whilst canonical TLR4-signalling strongly upregulates p62 and LC3, it sequesters these key elements of the autophagy machinery in the cytosol to drive inflammation in DCs by promoting p62-mediated NF-*κ*B activation^70^ and inducing blockage of the autophagic flux^71, 72^ that would otherwise act to limit inflammation^73^. By contrast, ES-62 induces low level and selective autophagic flux and autophagolysosomal degradation of key proinflammatory signal transducers, harnessing this homeostatic regulatory network to dynamically suppress hyper-inflammatory (IL-6, IL-12p70 and TNF*α*) responses in these cells^69^. Overall, whilst our pathway analysis of the functional classes of fully hypermethylated CpG sites across the various gene regions highlighted that exposure ES-62 *in vivo* appeared to act to reverse CIA-induced hypomethylation of the TSS regions of a range of genes involved in reprogramming of differentiation (Fig. 7d-f), it generally resulted in SFs displaying stable profiles of DNA hypermethylation quite distinct to those of both naïve- and CIA-SFs across the promoter, 1^st^ exon and gene body regions of genes (Fig. 7a-l).

The differential nature of the observed rewiring of SF methylome status was further underlined by the binary methylation gene signatures exhibited by naïve, CIA and ES-62-CIA SFs, revealed by heatmap analysis of genes displaying either 0 or >90% CpG methylation at the various gene regions in one or more of the treatment groups (Fig. 8a-d): the gene classes/functional responses differentially regulated in this binary manner in the three treatment groups are represented in the pie-charts for each of the gene regions (Fig. 8e). Cross-mining of these data with genes already implicated in the (SF) pathogenesis of RA (Supplementary Table 1) indicates that the resultant methylation signatures are physiologically relevant and reflect the differential protective and pathogenic phenotypes of the treatment groups. Critically, these data clearly confirm that ES-62 does not simply prevent pathogenic epigenetic remodelling of SFs during CIA but rather, induces further rewiring to a phenotype that is associated with suppression of inflammation and bone destruction in the joint.

**Fig. 8.**
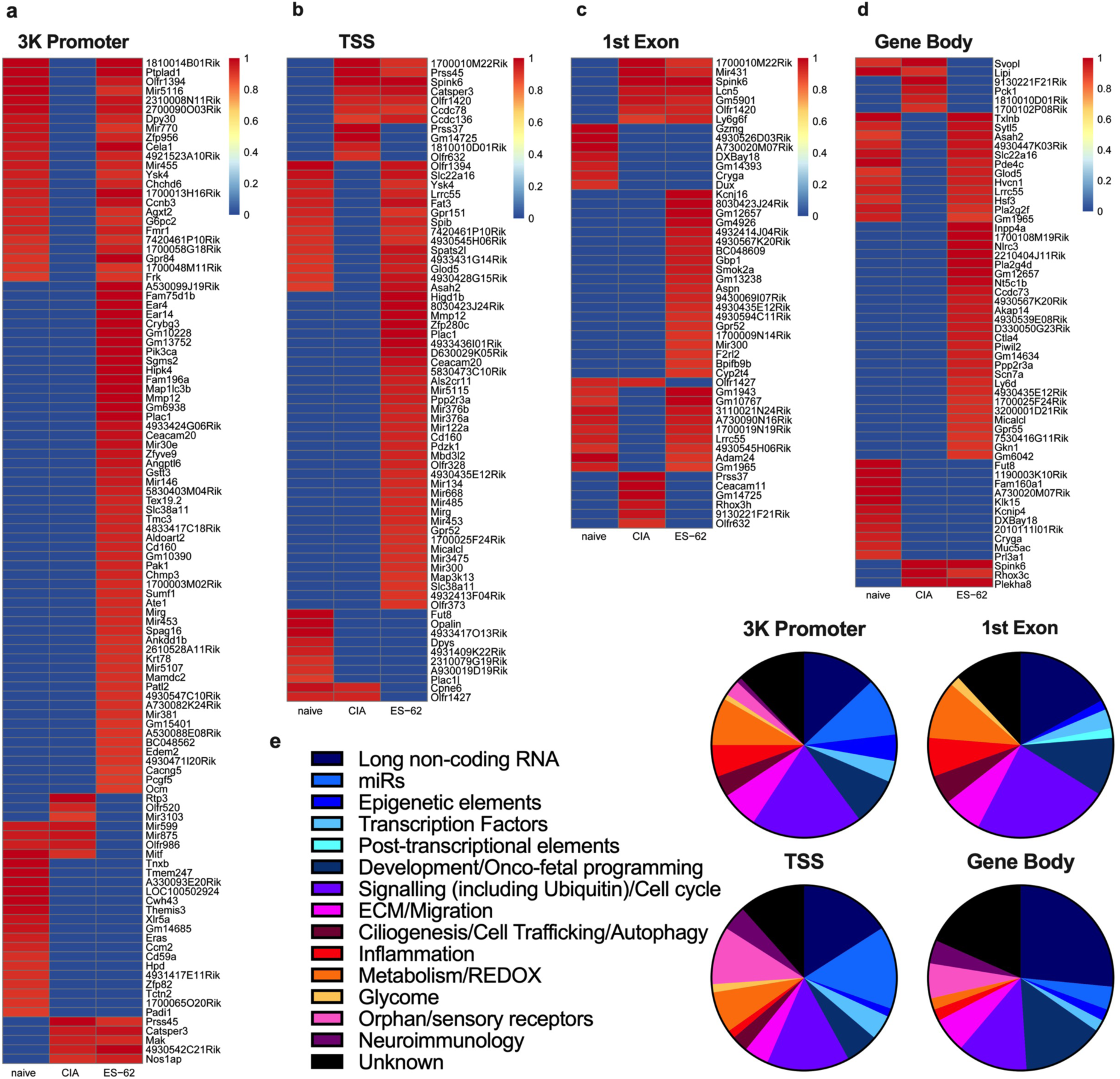
Binary analysis of the DNA methylome indicates differential silencing and activation of genes in SFs from Naïve, CIA- and ES-62 CIA-mice. Hierarchical clustering heatmaps were constructed for each of the (**a**) 3K Promoter, (**b**) TSS, (**c**) 1^st^ Exon and (**d**) Gene body regions analysing genes which were either essentially fully methylated (>0.9) or demethylated (0) at these sites in one or more of the treatment groups. (**e**) The functional classification of genes constituting these binary DNA methylation signatures differentially targeted in pathogenic (CIA) and protective (ES-62) rewiring of SFs are summarized in the pie-charts according to the accompanying colour key as indicated for each of the 3K Promoter, TSS, 1^st^ Exon and Gene body regions.

## Discussion

Collectively, our studies shed mechanistic light on how both acute and chronic interactions within the inflammatory microenvironment of the joint during induction and progression of CIA shape SF responses, resulting in remodelling of the epigenetic landscape in terms of DNA methylation and reprogramming to a hyperinflammatory, joint destructive cell phenotype (Fig. 9). This is a step-wise process characterised by increased expression of cytokines and growth factors (and their receptors), signalling molecules (e.g. Wnt, Ras-MAPK, FoxO pathways), adhesion molecules and extracellular matrix components and proteases^12–15^. Reflecting their high levels of expression in the CIA joint^17, 18, 74^, IL-1*β* and IL-17 are implicated in driving such reprogramming via DNA hypomethylation induced by downregulation of DNMT1 and evidenced by its recapitulation *in vitro*, following chronic exposure of naïve SFs to these pro-inflammatory factors and the DNMT1 inhibitor 5-azacytidine (5-aza). Moreover, reflecting a convergent upstream signalling pathway, both the acute (cytokine and MMP production) and chronic (remodelling to a stable aggressive phenotype) responses to these pathogenic cytokines are dependent on ERK and STAT3 signalling. Consistent with our previous studies demonstrating that the protection afforded by ES-62 against CIA was associated with the suppression of the aggressive phenotype of CIA-SFs^18^, we found the acute production of cytokines and MMPs by this stably imprinted “resolving” phenotype to be reduced to levels more typical of those of naïve SFs and this hyposensitivity was associated with a corresponding suppression of ERK and STAT3 signalling. Perhaps surprisingly therefore, these protective actions were not due to ES-62 simply preventing the transformation of SFs to the aggressive pathogenic phenotype but rather, reflected further epigenetic remodelling associated with a distinct phenotype that promotes resolution of inflammation and/or joint destruction (Fig. 9). Thus, given the functional differential outcomes of methylation of various gene sites^56–59^ and our finding that ES-62’s effects reflect even more profound global DNA hypomethylation than that observed in CIA-SFs, these studies suggest that utilizing interpretation of the global DNA methylation status of RA patients as a biomarker of disease is a strategy that should be treated with caution.

**Fig. 9.**
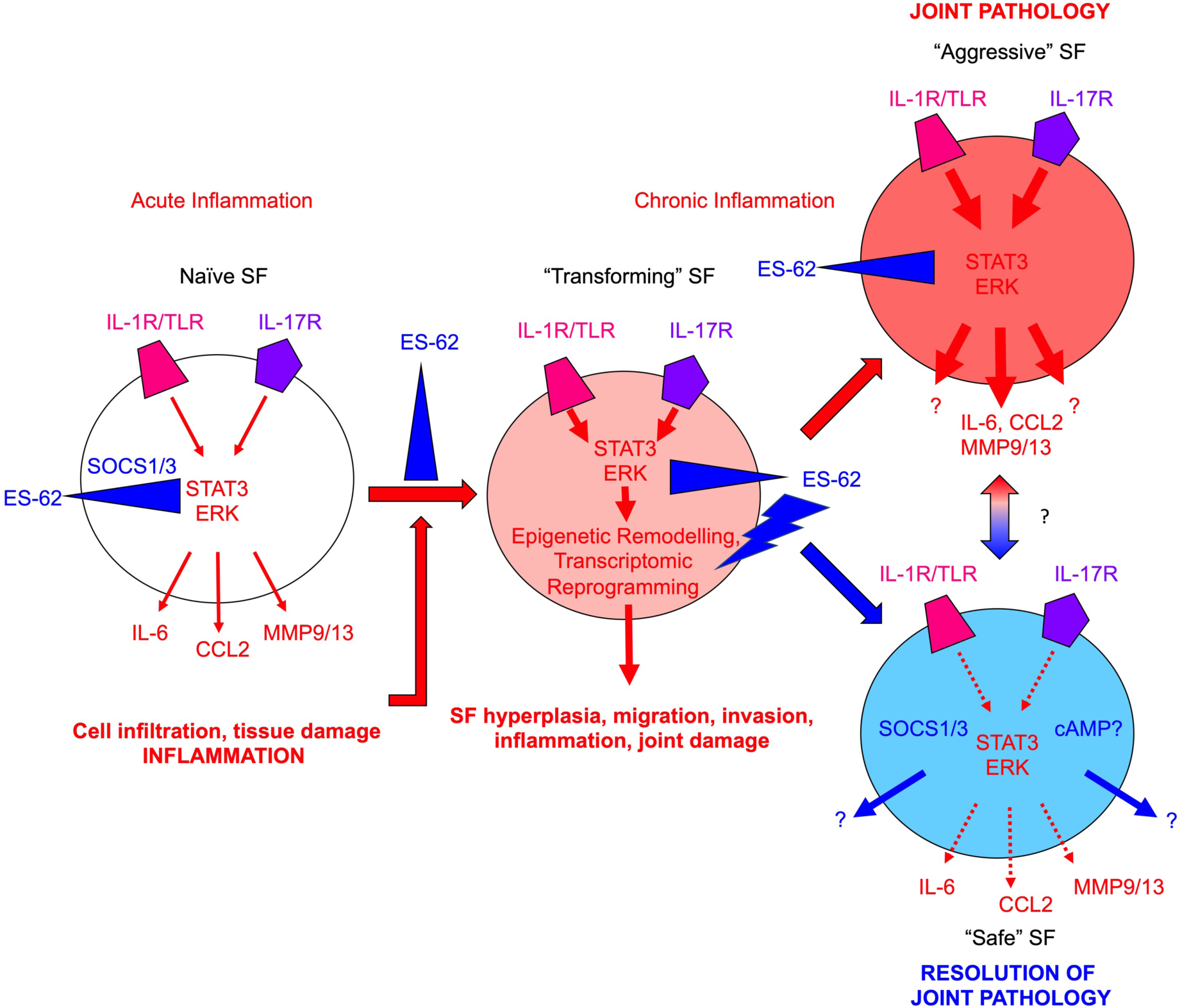
Model of ES-62 action in SFs during CIA. Healthy SFs (Naïve) respond to acute pro-inflammatory signals (e.g. IL-1*β*/IL-17) in the joint microenvironment resulting in release of pathological mediators such as IL-6, CCL2 and MMP9/13, which cause cell (neutrophils, macrophages, lymphocytes) infiltration and tissue damage, resulting in chronic inflammation. In addition to perpetuating joint inflammation and damage, chronic inflammation drives rewiring of the epigenetic landscape and consequently, reprogramming of the SFs to an “aggressive” hyperplastic, migrating and invasive phenotype that, via its hyper-responsiveness to environmental cues, plays a key role in joint pathology and destruction (all pathogenic signals denoted by red arrows with imprinted hyper-responsiveness represented by increased arrow size and redness of SF). ES-62 can disrupt this process at multiple points: thus, it can inhibit (blue blocking symbol) the (i) initial pro-inflammatory signalling to suppress acute inflammation; (ii) inflammatory signals generated by infiltrating cells and tissue damage to suppress induction of chronic inflammation; (iii) signalling associated with chronic inflammation to suppress transformation of SFs to an aggressive phenotype and (iv) pathogenic signals in aggressive SFs during established disease. Consistent with our findings that ES-62 does not simply maintain/restore the naïve SF phenotype, it also exhibits positive actions (blue lightning bolt symbol), specifically inducing epigenetic remodelling that reprograms the cell to an anti-inflammatory and tissue repair phenotype. The potential ability of the ES-62-CIA “safe” SFs to also directly influence the behavior of “aggressive” SFs in the joint during established disease is represented by the red-blue gradient arrow.

The ability of IL-1*β* and IL-17 to drive key features of SF pathogenicity^38, 75, 76^, and their targeting by ES-62^17, 18, 74^, underscores the importance of fully understanding mechanism of action in order to realise the potential of these cytokines (and/or their downstream signalling pathways) as candidate sites of intervention for therapeutically promoting re-establishment of joint homeostasis. Although the processes by which ES-62 suppresses ERK and STAT3 signalling in CIA-SFs have not been fully determined, we have previously shown the worm product to suppress ERK by both uncoupling upstream regulators and promoting MAPK phosphatase activity in innate and adaptive immune system cells^77^: consistent with this dual-pronged mechanism of action, our methylome analysis now suggests that ES-62 likewise acts to counter-regulate genes (e.g. Ysk4 [MAP3K19], Dusp1 [MAPK PTPase]) that could promote ERK signalling in SFs during CIA. Moreover, we have observed that ES-62 upregulates the important negative feedback regulators of inflammatory signalling, SOCS1 and SOCS3, that are also likely to impact on these pathogenic pathways in SFs. Thus, for example, SOCS1 can suppress IL-6 (indirect effector of IL-1*β*) and IL-1R/TLR (via targeting of Mal, IRAK1, Traf6 and NF-*κ*Bp65) signalling^22^ and both SOCS elements can directly negatively regulate JAK-STAT signalling, with typically (IFN-driven) STAT1 signalling being inhibited by SOCS1^78^ and (IL-1*β−*, IL-6- and IL-17-driven) STAT3 signalling suppressed by SOCS3^24^. Of relevance, miR-155 has been shown to negatively target SOCS1^22^, promoting the pathogenic IFN*γ* signalling^78–80^ in CIA that is targeted by ES-62^81, 82^. Similarly, miR- 146, which is found at elevated levels in RA-SFs and in response to key pathogenic mediators (IL-1*β*, IL-6, IL-17 and TNF-*α*), upregulates IRAK1 and Traf6 to enhance inflammatory signalling^38, 39^. Interestingly therefore, while we find miR-146 to be upregulated in CIA-SFs, this is not the case for these cells from ES-62-treated mice: such differential modulation of its expression is reflected at the methylome level with hypomethylation of the promoter region in CIA- but not ES-62-CIA SFs, suggesting that this miR may be an important target in the parasitic worm product’s rewiring of CIA-SFs. Moreover, we find that miR-455 (Supplementary Table 2), which acts to downregulate SOCS3^83^ is likely induced (hypomethylated at its promoter 3K region) in CIA SFs but silenced (hypermethylated promoter) in naïve and ES-62-CIA SFs. Thus, the finding that ES-62 reduces miR-146, -155 and -455 levels suggests that this might provide an acute and dynamic mechanism by which it maintains levels of SOCS1 and SOCS3 during CIA to regulate and resolve pathogenic SF responses elicited by IL-1R/TLR and cytokine signals chronically generated in the inflamed joints.

Methylome analysis of potential targets underpinning the transformation of SFs to the aggressive dedifferentiated/tumor-like phenotype characterized by hyperplasia and aberrant migration now sign-posts other, new, candidate pathways for exploration of ES-62 action and potential therapeutic intervention. Thus, whilst naïve SFs display hypermethylation of the promoter and TSS regions of genes associated with regulation of pluripotency and (oncofetal) development, many of these sites are hypomethylated and potentially derepressed in CIA-, but not ES-62-CIA-, SFs. By contrast, whilst elements associated with the FoxO, Hippo and Wnt signalling pathways that are key to the regulation of these processes (stem cell biology, cilia and intercellular communication, organ development, immunomodulation, tissue repair and regeneration and ageing^64, 84, 85^) are similarly hypomethylated in CIA SFs and hypermethylated in ES-62-CIA, these genes are also hypomethylated in naïve SFs. Of related interest, we have previously shown ES-62 to strongly induce negative regulators (WIF1, AMOTL2 and CRYBB2) of Wnt signalling in RA synovial membranes, *in vitro*^86^. Critically, this silencing (hypermethylation) of genes associated with ciliogenesis and its regulation of signalling and differentiation is not a reversion to the naive SF phenotype, underscoring that ES-62 drives rewiring of a unique and “safe/protective” phenotype of SF in mice undergoing CIA. Similarly, whilst induction of CIA is associated with hypermethylation (silencing) of the promoter region of Taf9b (coactivator and stabilizer of p53 and negatively regulated by miR146^87^), the promoter and TSS sites respectively of Ccnb3 (G2/mitosis cyclin B3) and MAP3K13 (stabilizes c-Myc to promote survival, proliferation and tumorigenesis^88^) are hypomethylated: ES-62 inverts the methylation status of all 3 of these genes, reverting to the naïve status of Taf9b and Ccnb3 and the ES-62-CIA-specific phenotype for MAP3K13. Moreover, with respect to the aberrant migration of SFs, the promoter region of MAPK10, a signalling element whose downregulation is associated with migration, invasion and angiogenesis in cancer^89^ is hypermethylated in CIA- but not naïve or ES-62-CIA, SFs. At the same time, exposure to ES-62 uniquely hypermethylates the promoters of genes implicated in ciliogenesis (MapKap1 [Sin1 subunit of mTORC2], Inpp5b, DynII2, Chmp3, Pik3ca) and migration and interaction with the ECM (Cela1 [chymotrypsin-like elastase family member 1], Ate-1, Spag16). Collectively therefore, whilst CIA-induced methylation changes may contribute to the functional reprogramming underpinning the observed hyperplasia, migration and hyper-inflammatory SF responses, ES-62 appears to induce a safer SF phenotype involving both prevention of a number of these changes but also further epigenetic rewiring of additional genes.

Support for the translational relevance of the differential DNA methylation signatures of SFs associated with naïve, CIA and ES-62-CIA mice is provided by our cross-data mining of genes implicated in RA pathogenesis which highlighted that our observed differential epigenetic regulation of key factors (e.g. *miR-146, Runx3, Batf, Pak1, Pde4c, Mmp12, Spag16* and various cytokines, chemokines and immunomodulatory receptors) was functionally predictive of both CIA pathogenesis and ES-62-mediated suppression and resolution of joint inflammation and destruction. Perhaps the observed counter-regulation of *Pde4c* is particularly pertinent to current translational strategies given the recent strong interest in the therapeutic potential of PDE4 inhibitors in RA as well as a wide range of other chronic inflammatory conditions including psoriatic arthritis, for which Apremilast was recently approved as a treatment^90^: indeed, ES-62 further acts to promote cAMP signalling in SFs by inducing inverse methylation signatures in the 1^st^ Exon region of other important elements, including Adcy10 (adenylate cyclase 10), AKAPs (A-kinase anchoring proteins) and Prkacb (Protein kinase A catalytic subunit beta) to those of naïve- and CIA-SFs. Interestingly therefore, in the context of ES-62 targeting this anti-inflammatory pathway, cilia provide a spatiotemporal platform for compartmentalising elements of the cAMP signalsome to promote protein kinase A-mediated antagonism of Hedgehog signalling^91^, a key pathway regulating cell differentiation which has recently been implicated in the proliferation, migration and invasion of RA SFs^92^. Moreover, recent mathematical modelling of differential gene expression metadata from a number of studies in healthy and RA synovial tissue^93^ identified a gene signature panel representing an interactive biological network of hub (STAT1, RAC2 and KYNU) proteins and effector molecules (PEPD receptor and NR4A1, MEOX2, KLF4, IRF1 and MYB transcription factors and miRs-146a, -299, -3659, -6882 and - 8078) that provides a platform for validation of pathogenic mechanisms and potentially, a predictive tool for diagnosis and development of effective treatment strategies for RA: apart from KYNU and miR-299 (miRs-3659, -6882 and -8078 were not identified in our methylome study), these “signature” genes were differentially methylated at one or more sites in naïve-, CIA- and ES-62-CIA-SFs (Supplementary Information Table 2), with CIA- and ES-62 SFs generally exhibiting inverse profiles of methylation.

Collectively, our data suggest that in addition to acutely suppressing pathogenic SF responses, ES-62 acts to stably reset homeostatic regulation of inflammation-resolving and tissue repair genes by epigenetically rewiring these cells away from their transformed aggressive phenotype in the arthritic joint. Importantly, the finding that ES-62 does not simply prevent generation of the pathogenic phenotype but rewires it to a stably protective form should have translational impact for treating established joint disease. Interestingly, this reprogramming of SFs is not likely to be the only instance of such epigenetic rewiring activity by the parasite product as although not examined at the level of the methylome, we have previously shown ES-62 to induce stable anti-inflammatory phenotypes of dendritic cells (DCs) and macrophages (from bone marrow-derived progenitors) and immunoregulatory B cells in mouse models (CIA and MRL/Lpr mice) of RA and SLE^1, 94, 95^. Interestingly therefore, the suppression of DC maturation (buffalo cells), particularly in terms of TLR signalling, by excretory-secretory products released by the parasitic trematode worm *Fasciola gigantica* (FgESPs) has recently been shown to be associated with changes in the methylome and transcriptome reprogramming^96^. Moreover, the methyl-CpG-binding protein, Mbd2, that acts to coordinate chromatin accessibility and hence, reprogramming of gene expression in response to DNA methylation, is key to the ability of DCs to drive Th2 responses to helminths and allergy^97^ and also plays a role in CD11c+ DC/monocyte-epithelial cell crosstalk in gut inflammation^98^. Although, exposure to ES-62 does not substantially impact on the methylation profile of Mbd2 in CIA-SFs, it results in TSS hypermethylation of Mbd3l2, a homologous protein that can also interact with components of the NuRD complex but to antagonize Mbd2-mediated methylation silencing^99, 100^, an inverse signature to that of naïve and CIA-SFs (hypomethylated). Thus, such protective rewiring of both haemopoietic and stromal cells is perhaps an underexplored feature of the co-evolution of the host-parasitic worm relationship to date. However, taken in the context of our recent findings that ES-62 can promote both health- and life-span in male mice undergoing obesity-accelerated ageing^6^, it is in line with helminths having evolved such an overarching inflammation-resolving and repair/regeneration strategy to counter the chronic inflammation and tissue pathology they would otherwise elicit. Consistent with this, it has recently been reported that another parasitic trematode, *Schistosoma mansoni,* induces hepatocyte DNA hypomethylation that is associated with reduced tissue pathology and granuloma formation in the acute phase of infection in mice^101^, supporting the idea that epigenetic changes induced by helminth infection appear to be indirectly important in promoting long-term helminth survival.

Interestingly, such epigenetic remodelling was impacted by the inflammatory context, which modulated the (reciprocal) epigenetic changes occurring in both the host hepatocytes and the worms to determine the levels of liver pathology^101^. The potential importance of the evolution of such complex host-worm epigenetic regulatory networks is highlighted by evidence that CD4 T cells, from children recently exposed to tuberculosis and infected with *Schistosoma haematobium* showed profound differential DNA methylation, relative to those from uninfected children, that was associated with Th2-skewing of tuberculosis (TB)-specific responses away from the protective Th1/IFN*γ* phenotype^102^. Moreover, the remodelled DNA-methylation signatures and defective TB-specific Th1 responses were maintained for up to at least 6 months following successful deworming of *S. haematobium*^102^. Furthermore, intriguingly, emerging evidence suggests that the offspring of chronically-infected mothers display epigenetically rewired immune responses that potentially shape their reactions to other pathogens (bacterial and viral infections) and vaccines^103, 104^. In conclusion therefore, understanding these interactive epigenetic networks has important implications not only for how to develop strategies to counter the rise in prevalence of chronic inflammatory (allergic and autoimmune) conditions, increasingly associated with eradication of helminths, but also to combat pathogens in a co-infection setting.

## Materials and Methods

### Collagen-induced arthritis (CIA)

DBA/1 (Envigo) mice were maintained in the Central Research Facility of the University of Glasgow and all studies were approved by and performed in accordance with UK Home Office regulations and guidelines (Licences PPL60/26532, I675F0C46, PIL70/26532) and the Animal Welfare and Ethics Review Board of the University of Glasgow. For the CIA model, male DBA/1 mice (8-weeks old) were injected intradermally on day 0 with 100 µg Bovine Type II collagen (CII; MD biosciences) in complete Freunds adjuvant (CFA; MD biosciences) just above the tail base. On Day 21, mice were administered 200 µg of type II collagen in PBS via intra-peritoneal injection^105^. Mice were monitored every two days for signs of arthritis as described previously, being determined according to the following articular scores: 0 = normal; 1 = digit(s) involvement; 2 = erythema; 3 = erythema and swelling; 4 = extension/loss of function and in addition, by calliper measurement of hind paw swelling (POCO 2T, Kroeplin Längenmesstechnik). Where indicated, ES-62 (2 µg in PBS, prepared endotoxin-free and purified as previously described^105^) was administered subcutaneously in the scruff on days -2, 0 and 21.

SFs were isolated and cultured according to the protocol of Armaka *et al* as described previously^106^. Briefly, crushed joints were treated with 1 mg/ml collagenase IV (*Clostridium histolyticum*; Sigma) at 37°C for 1 hour and the released cells resuspended in DMEM (Lonza), supplemented with 10% FBS, 2mM L-glutamine, and 50 units/ml penicillin and 50 µg/ml streptomycin, and plated in 100 mm dishes. After 24 hours, non-adherent cells were removed and the cells passaged (medium replaced twice weekly) and expanded for 3 to 4 weeks before analysis. The SF phenotype of the explant cultures (CD54^+^CD11b^-^CD90.2^+^) was confirmed by flow cytometry^18^.

Analysis of *in vivo* hypoxia status of SFs was determined using Pimonidazole HCl (dose of 60 mg/kg; Hypoxyprobe^TM_^1 plus kit protocols) injected intra-peritoneally 24h prior to cull^107^. Joint cells were extracted and then fixed in chilled 70% ethanol and stored at -20°C for 24 hours. Cells were next resuspended in PST (PBS with 4% serum and 0.1 % [v/v] Triton X-100) and stained using the kit FITC-conjugated Mab1 (diluted 1:1000) for 2 hours at 37°C prior to analysis by flow cytometry. Likewise, the vasculature permeability occurring *in vivo* was assessed following intravenous injection (200 µl) of Evans blue dye (Sigma; 0.5% in PBS) in the mouse lateral tail 30 min prior to cull^108^. For histochemistry, following fixation and decalcification of bones, joints were paraffin-sectioned and stained with H&E or Trichrome solutions as described previously^18^.

### Flow Cytometry

SFs (0.5 x 10^6^) were stained with the fixable viability dye eFluor® 780 (1 μg/ml; eBioscience) prior to blocking Fcɣ receptors and incubation with the indicated primary (5 µg/ml anti-CD11b-FITC [M1/70]; 2 µg/ml anti-CD54-PE [YN1/1.7.4]; 2 µg/ml anti-CD90.2-PerCP [30H12] or appropriate isotype controls) and where relevant, secondary (streptavidin conjugated fluorophores) antibodies. For intracellular staining of signalling molecules, cells were first stimulated as indicated prior to permeabilization with methanol and then ERK protein (ERK), dually phosphorylated ERK (pERK), STAT-3 protein (STAT-3) or phosphorylated (pSTAT-3) expression detected by the relevant rabbit antibodies (all Cell Signalling Technology) and secondary FITC-conjugated anti-rabbit IgG antibodies (Biolegend) as described previously^109^. All samples were acquired using the BD FACSCalibur or LSR II flow cytometers and analysed by FlowJo software.

### Determination of cytokine release

SFs were seeded at 10^4^ (96-well plates; Corning) and 0.3 x10^6^ (6-well plates) cells per well and after overnight incubation in 1% FCS DMEM to synchronize the cells, stimulated for 24 hours in 10% FCS DMEM with LPS (1 µg/ml, *Salmonella Minnesota*; Sigma-Aldrich), BLP (Pam_3_Cys-Ser-(Lys)_4_; 0.5 µg/ml, Enzo), IL-1β (10 ng/ml, Immunotools) or IL-17 (25 ng/ml, Immunotools) as indicated. Supernatants were collected and analysed using ELISA kits detecting IL-6, CCL-2 and IL-1β (Ready-SET-Go!® from eBioscience), with absorbance read at 450nm using a Tecan Sunrise plate reader.

### FACE assays

Fast-activated cell-based (FACE) ELISAs were performed as we described previously^110^. SFs were seeded at 10^4^ cells/well (96-well plates) and after overnight incubation in 1% FCS DMEM to synchronize the cells, were then stimulated for the indicated time points (from 0 to 120 minutes) before being treated with fixative buffer (Biolegend) according to the manufacturer recommendations. Once permeabilized, the cells were incubated with antibodies specific for either the activated or non-activated form of the signalling element (e.g., ERK1/2 and the dually phosphorylated active forms of ERK [pERK; Cell Signalling Technology] for 1 hour before addition of streptavidin-conjugated horseradish peroxidase for 30 minutes (anti-rabbit HRP, Cell Signalling Technology). Following washing and incubation with tetramethylbenzidine (TMB), absorbance was read at 450nm using a Tecan Sunrise plate reader.

### Western Blot analysis

SFs were seeded in 6-well plates at 1×10^6^ cells per well and synchronised overnight in 1% FCS DMEM before being treated for 24 hours with the indicated stimuli: LPS (1 µg/ml, Sigma-Aldrich), IL-1β (10ng/ml, Immunotools) or IL-17 (25 ng/ml, Immunotools). Following washing, cells were lysed by addition of 100 µl of RIPA buffer (50 nM Tris buffer pH 7.4, 150 mM sodium chloride, 2% (v/v) NP40, 0.25% (w/v) sodium deoxycholate, 1 mM EGTA, 1x Halt^TM^ protease inhibitor and 1x Halt^TM^ phosphatase inhibitor [Pierce]) and the protein concentration of the solubilised proteins measured by the BCA protein assay kit (Pierce). Proteins (20 µg) were separated by SDS-PAGE on 4-12% Bis-Tris gradient gels, using the NuPAGE Novex system (Invitrogen) in NuPAGE MOPS buffer, with addition of antioxidant. Proteins were then transferred onto nitrocellulose membranes (GE) using the wet transfer NuPAGE system as described previously^69^. Following blocking (5% non-fat milk protein in Tris buffered saline-1% (v/v) Tween-20 [TBS-T]), membranes were incubated with the relevant primary antibody in 5% BSA TBS-T overnight at 4°C. After washing, secondary antibody conjugated to HRP was added in 5% milk for 1 hour at room temperature and following washing developed using ECL Western blotting substrate (Thermo Fisher Scientific) and X-ray film (Kodak). Membranes were stripped and re-probed using Restore Western Blot stripping buffer (Thermo Fisher Scientific) as described previously^69^.

### mRNA and miRNA analysis

Total RNA and miRNA were isolated and separated using the miRNeasy kit (Qiagen) following the manufacturer’s instruction.

For miRNAs, the miScript Reverse Transcription Kit (Qiagen) was used for cDNA preparation. miScript SYBR green qPCR kit (Qiagen) and miScript primer assay (Qiagen) were used for semi-quantitative determination of the expression of mouse miR-19b-1 (MS00005915), miR19b-1* (MS00024493), miR-19b-2* (MS00024500), miR-23b-2 (MS00032606), miR-24-1* (MS00011543), miR-34a (MS00001428), miR-124*(MS00011081), miR-125a (MS00001533) miR-146 (MS00001638), miR-155 (MS00001701), miR-203 (MS00001848), miR-203* (MS00011452) and miR-346_2 (MS00032753). The expression of Hs-RNU6-2_11 (MS00033740) was used as endogenous control^111^. SFs were transfected with the miR-155 mimic, miR-155-5p or its control scrambled miR (Thermo Scientific Dharmacon) using Effectene (Qiagen) according to the manufacturer’s instructions as described previously^112^.

For mRNAs, the high capacity cDNA reverse transcription kit (Invitrogen) was used and subsequently, qPCRs were performed using Taqman® gene expression assay kits, with all primers obtained from Life Technologies for the following genes: CCL2 (Mm00441242_m1), DNMT1 (Mm01151063_m1), DNMT3a (Mm00432881_m1), GAPDH (Mm03302249_g1), IL-6 (Mm00446190_m1), MMP9 (Mm00442991_m1) and MMP13 (Mm00439491_m1).

### DNA methylation analysis

Global DNA methylation levels were determined by ELISA-like measurement of methyl cytosine levels. Briefly, genomic DNA was extracted from cells (PureLink Genomic DNA Mini Kit, Invitrogen) and methyl cytosine levels assessed by a specific primary and secondary HRP-conjugated antibody system following the manufacturer’s instructions (Imprint® Methylated DNA Quantification Kit-Sigma). Following development with TMB, absorbance was read at 450nm using a Tecan Sunrise plate reader to determine relative global DNA methylation levels. In experiments investigating induced DNA-demethylation of naïve SFs following chronic exposure to cytokines, cells were first synchronized overnight in 1% FCS DMEM and then cultured for 14 days in 10% FCS DMEM and stimulated daily with IL-1β (1 ng/ml; Invitrogen) or IL-17 at (2.5 ng/ml; Invitrogen), following the protocol previously published for RASFs^113^. In addition, as a control for the global DNA methylation assay, following synchronisation, SFs were treated daily with the DNMT1 inhibitor, 5-azacytidine (5-aza, Sigma-Aldrich; 1 µM in 10% FCS DMEM) for 7 days according to previous publications^12, 46^.

DNA methylome analysis of SFs (10^6^ cells/group) pooled from paws representative of the naïve (n=6), CIA (articular score, 3.17 ± 1.38, n=6) and ES-62-CIA (articular score, 0.5 ± 0.22, n=6) cohorts, from DNA purification through Illumina sequencing to bioinformatics, was by the commercial Active Motif Reduced Representation Bisulfite Sequencing (RRBS) single base pair resolution DNA methylation and bioinformatic service. Quantifying methylation for individual CpG motifs relies on using multiple sequencing reads to calculate methylation frequencies and as there are always tags sequenced that are a result of PCR duplication, replicates may bias the data analysis. Rather, specificity is provided by “barcoding” a random hexamer into the library sequencing adaptors that serves as a unique signature for each DNA fragment, distinguishing distinctive sequencing tags from tags generated by qPCR duplication and allowing greater data (>4m CpGs, with minimum sequence coverage of 3) accuracy and retrospective individual analysis despite reducing the required sequencing depth from 10 to 3. Thus, analysis was performed on pooled samples from mice of similar disease score within a treatment group basis in order to minimise variation arising from the factor we could control for, namely differences in the joint pathology arising in individual mice, a strategy we have previously successfully employed to investigate the differential metagenomic signatures of ES-62-treated mice undergoing CIA and obesity-accelerated ageing^6, 16^. Briefly, genomic DNA was extracted using the Quick-gDNA MiniPrep kit (Zymo Research D3024) following the manufacturer’s instructions for working with cell suspensions and proteinase K digested samples. gDNA (100 ng) was digested with TaqaI (NEB R0149) at 65⁰C for 2h followed by MspI (NEB R0106) at 37⁰C overnight. Following enzymatic digestion, samples were used for library generation using the Ovation RRBS Methyl-Seq System (Tecan 0353-32) following the manufacturer’s instructions. In brief, digested DNA was randomly ligated, and, following fragment end repair, bisulfite converted using the EpiTect Fast DNA Bisulfite Kit (Qiagen 59824) following the manufacturer’s protocol. After conversion and clean-up, samples were amplified resuming the Ovation RRBS Methyl-Seq System protocol for library amplification and purification. Libraries were measured using Agilent 2200 TapeStation System and quantified using the KAPA Library Quant Kit ABI Prism qPCR Mix (Roche KK4835). Libraries were sequenced on a NextSeq 500 at SE75 and the reads mapped to the genome using RRBSMAP and PCR duplicates removed by proprietary script: the general sequencing statistics are shown in Supplementary Information, Table 3. Data were visualised by uploading BED files within the UCSC genome browser.

For pathway analysis, genes were filtered according to their methylation content in the 3K promoter, TSS, 1st Exon and gene body regions. The following filter conditions were selected to identify “binary” methylation signatures of synovial fibroblasts: (i) naïve ≥ 0.9, CIA ≥ 0.9, ES-62 = 0; (ii) naïve = 0, CIA ≥ 0.9, ES-62 ≥ 0.9; (iii) naïve ≥ 0.9, CIA = 0, ES-62 ≥ 0.9; (iv) naïve ≥0.9, CIA=0, ES-62=0; (v) naïve = 0, CIA ≥ 0.9, ES-62 =0 and (vi) naïve = 0, CIA = 0, ES-62 ≥ 0.9. Values of 1 represent full methylation of selected region whilst 0 represents no methylation found. Heatmaps to show these “binary” differential methylation profiles were made using the heatmap function under R environment. Each set of genes (methylation content ≥ 0.9 for all three treatments) were further uploaded into String software (http://string-db.org, ELIXIR) to identify significantly regulated KEGG pathways in interactive gene clusters and functional gene classifications (summarised by pie-charts).

### Statistics

All statistical analyses were performed using Prism software (GraphPad Software) using the one-tailed Student’s t-test (parametric data) or Mann-Whitney test (non-parametric data) and One- or Two-way ANOVA with appropriate post-test as indicated. Statistical significance is shown as *p<0.05, **p<0.01 and ***p<0.001.

## Data Availability

The data are contained within the article and Supplementary Information files. On publication, further information regarding experimental design and the DNA methylation data will be accessible via the linked Research Reporting summary and relevant accession numbers to be provided in the Method section, respectively. All other primary data files will be available from the corresponding authors on reasonable request.

## Acknowledgements

The study was funded by awards from the Wellcome Trust (086852) and Arthritis Research UK (21133) to M.M.H and W.H. M.A.P. is a Versus Arthritis Fellow (21221) and M.C. was awarded a Ph D studentship from the Wellcome Trust.

## Author contributions

M.C., M.A.P., A.T. and S.McG. conducted the experiments for the study which was designed by M.C., M.M.H. and W.H. R.N. provided the tissue from miR-155-LacZ reporter strain of mice and F.E.L. maintained the *A. viteae* life cycle and purified ES-62. M.A.P., K.Y. and A.T. performed the bioinformatics pathway analysis of the DNA methylome data. M.M.H. and W.H. wrote the paper, with all authors contributing their advice and being involved in reviewing and revising the manuscript and approving the final version.

## Supplementary Information

**Supplementary Table 1:**
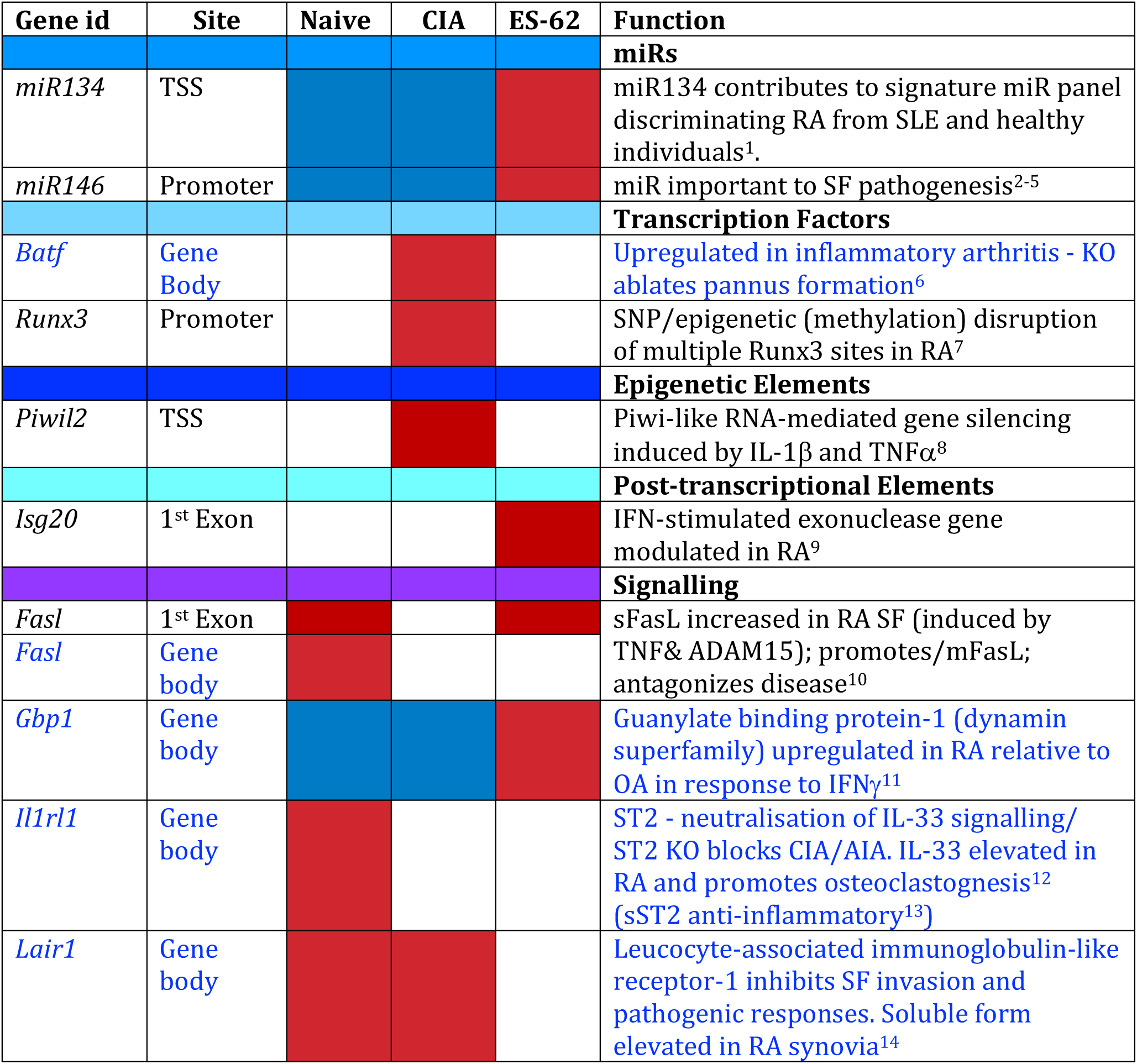
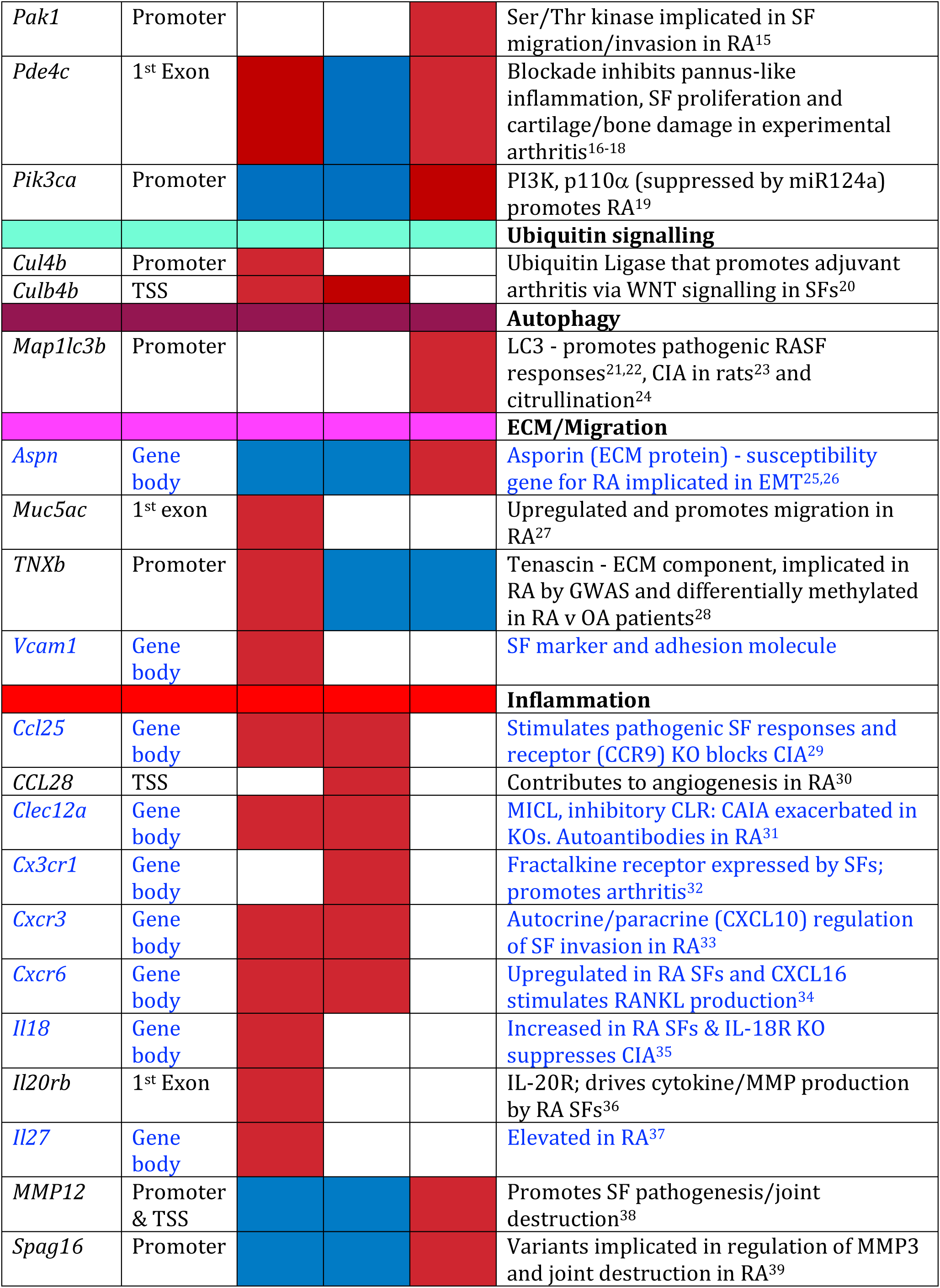
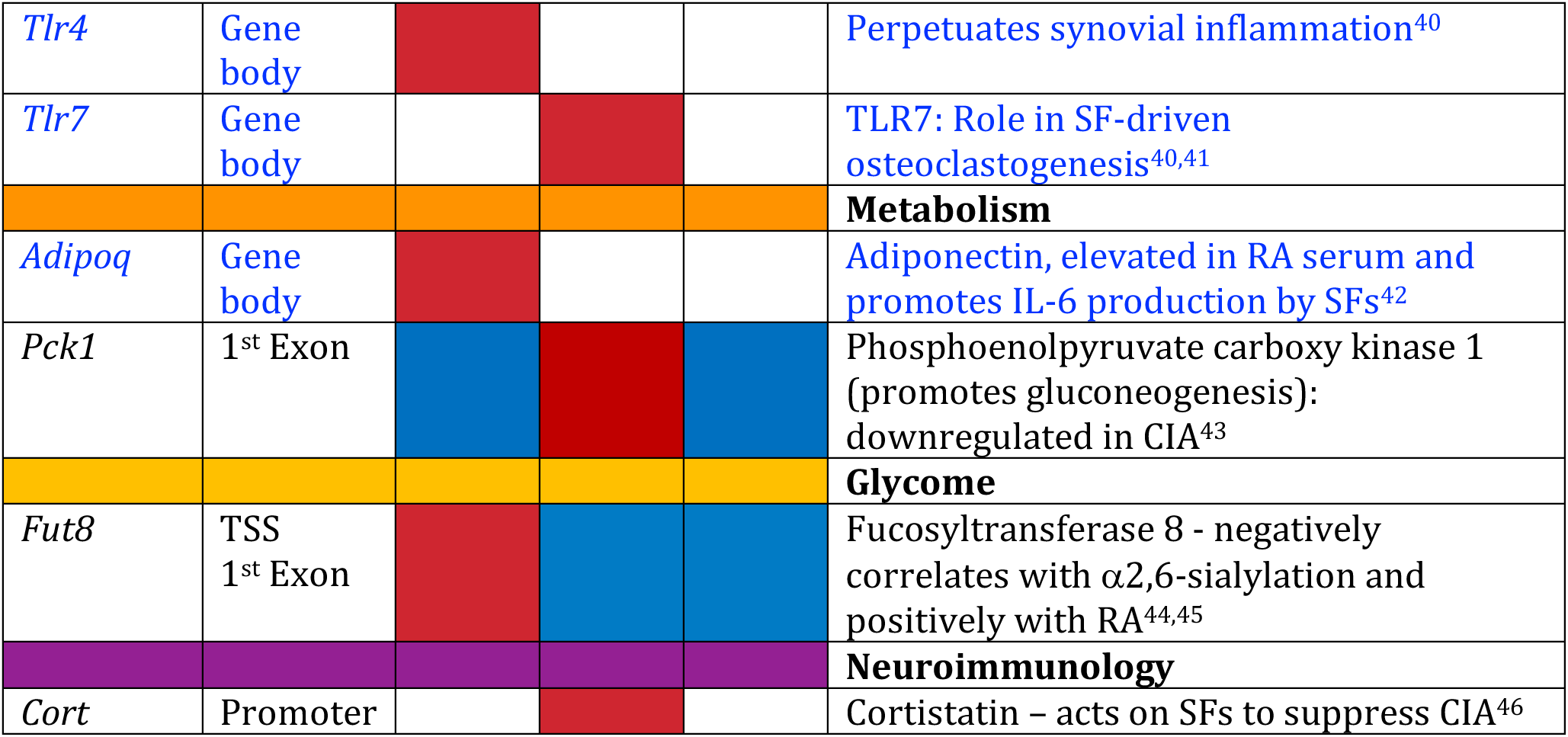
**Differential expression of genes implicated in Rheumatoid Arthritis Synovial Fibroblast pathogenesis.** The classes of genes targeted are colour-coded according to the pie-charts in Figures 7 and 8, whilst the treatment groups are coloured brick red and blue to reflect the differential methylation (Fig. 8) at the particular sites, highlighted in the ***heatmaps***. In addition, genes in clusters identified by the ***pathway analysis*** (Fig. 7) are coloured as brick red for >90% methylation whilst white indicates 0< methylation <90%. Genes targeted in the gene body region are denoted in blue font.

---

**Supplementary Table 2:**
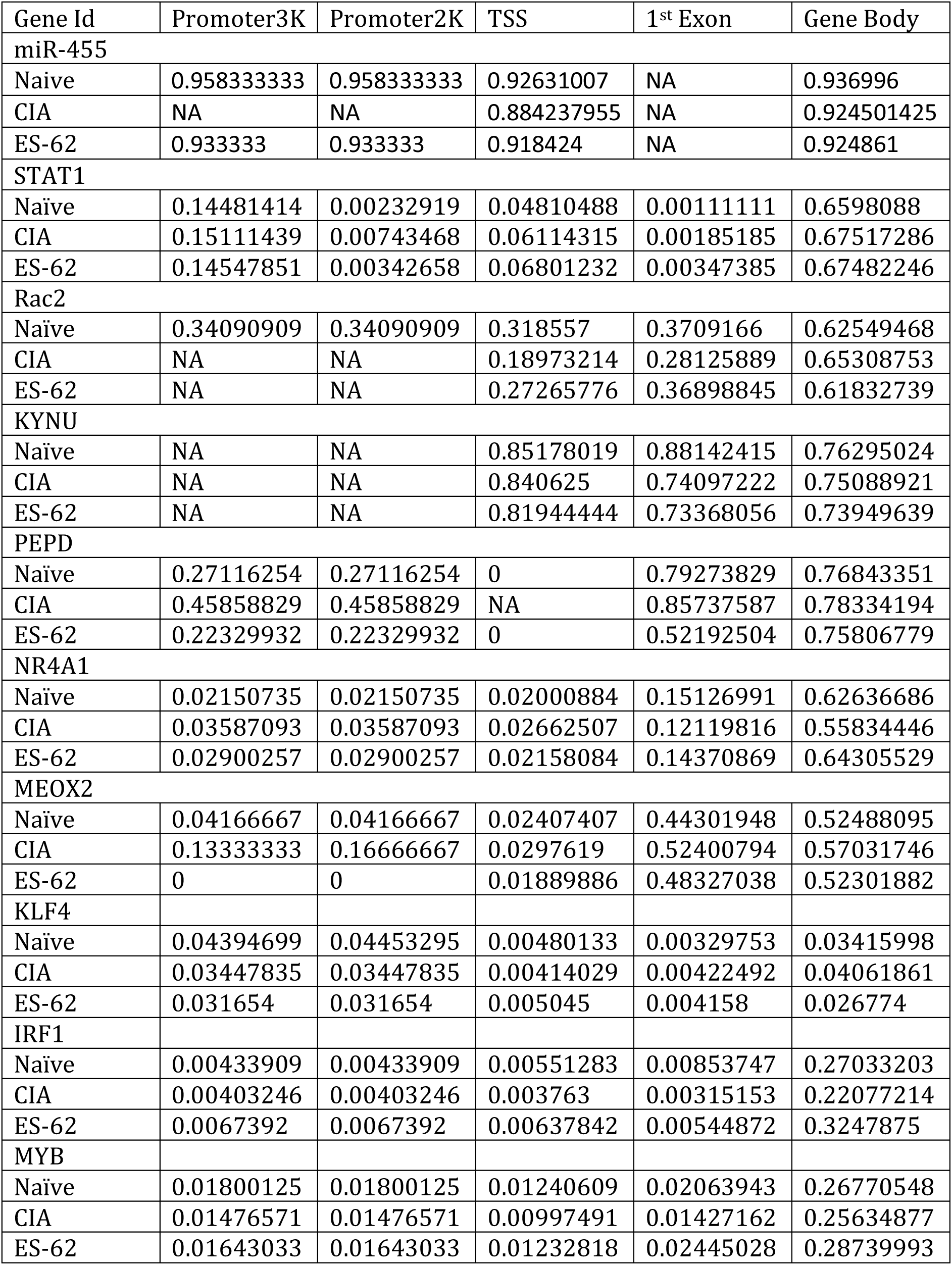
Methylation Status (where 1 represents fully methylated) of Genes identified as RA synovial signature

**Supplementary Table 3.**
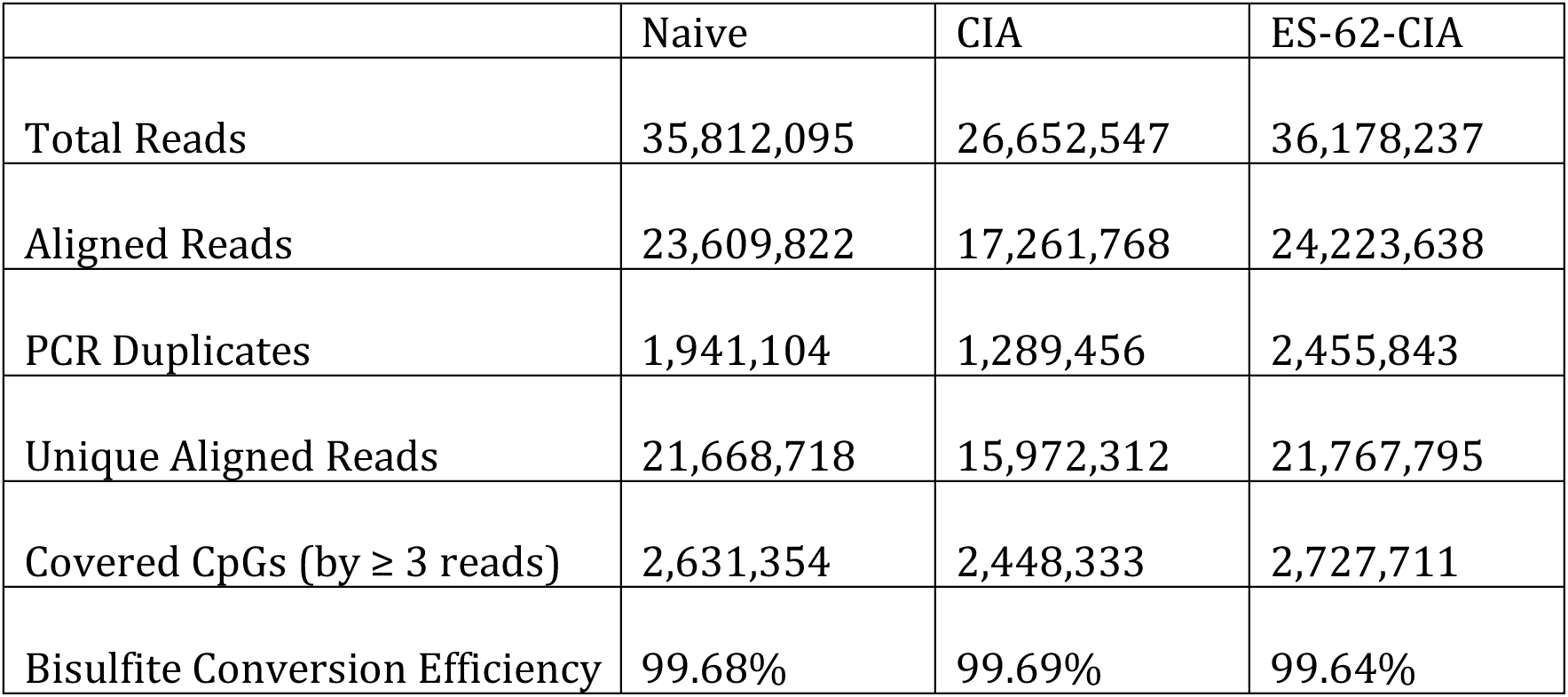
Sequencing Statistics of RRBS analysis

|  | Naive | CIA | ES-62-CIA |
| --- | --- | --- | --- |
| Total Reads | 35,812,095 | 26,652,547 | 36,178,237 |
| Aligned Reads | 23,609,822 | 17,261,768 | 24,223,638 |
| PCR Duplicates | 1,941,104 | 1,289,456 | 2,455,843 |
| Unique Aligned Reads | 21,668,718 | 15,972,312 | 21,767,795 |
| Covered CpGs (by $\geq 3$ reads) | 2,631,354 | 2,448,333 | 2,727,711 |
| Bisulfite Conversion Efficiency | 99.68% | 99.69% | 99.64% |

## References

1 Harnett, M. M. & Harnett, W. Can Parasitic Worms Cure the Modern World’s Ills? Trends in Parasitology 33, 694–705, doi:10.1016/j.pt.2017.05.007 (2017).

2 Panda, A. K. & Das, B. K. Absence of filarial infection in patients of systemic lupus erythematosus (SLE) in filarial endemic area: a possible protective role. Lupus 23, 1553–1554, doi:10.1177/0961203314546019 (2014).

3 Panda, A. K. & Das, B. K. Diminished IL-17A levels may protect filarial-infected individuals from development of rheumatoid arthritis and systemic lupus erythematosus. Lupus 26, 348–354, doi:10.1177/0961203316662722 (2017).

4 Panda, A. K., Ravindran, B. & Das, B. K. Rheumatoid arthritis patients are free of filarial infection in an area where filariasis is endemic: comment on the article by Pineda et al. Arthritis and Rheumatism 65, 1402–1403, doi:10.1002/art.37883 (2013).

5 Harnett, W. Secretory products of helminth parasites as immunomodulators. Molecular and Biochemical Parasitology 195, 130–136, doi:10.1016/j.molbiopara.2014.03.007 (2014).

6 Crowe, J. et al. The parasitic worm product ES-62 promotes health- and life-span in a high calorie diet-accelerated mouse model of ageing. PLoS Pathogens 16, e1008391, doi:10.1371/journal.ppat.1008391 (2020).

7 Juarez, M., Filer, A. & Buckley, C. D. Fibroblasts as therapeutic targets in rheumatoid arthritis and cancer. Swiss Med Wkly 142, w13529, doi:10.4414/smw.2012.13529 smw-13529 [pii] (2012).

8 Nygaard, G. & Firestein, G. S. Restoring synovial homeostasis in rheumatoid arthritis by targeting fibroblast-like synoviocytes. Nat Rev Rheumatol 16, 316–333, doi:10.1038/s41584-020-0413-5 (2020).

9 Haniffa, M. A., Collin, M. P., Buckley, C. D. & Dazzi, F. Mesenchymal stem cells: the fibroblasts’ new clothes? Haematologica 94, 258–263, doi:10.3324/haematol.13699 (2009).

10 Steenvoorden, M. M. et al. Transition of healthy to diseased synovial tissue in rheumatoid arthritis is associated with gain of mesenchymal/fibrotic characteristics. Arthritis Research & Therapy 8, R165, doi:10.1186/ar2073 (2006).

11 Zvaifler, N. J. Relevance of the stroma and epithelial-mesenchymal transition (EMT) for the rheumatic diseases. Arthritis Research & Therapy 8, 210, doi:10.1186/ar1963 (2006).

12 Karouzakis, E., Gay, R. E., Michel, B. A., Gay, S. & Neidhart, M. DNA hypomethylation in rheumatoid arthritis synovial fibroblasts. Arthritis and Rheumatism 60, 3613–3622, doi:10.1002/art.25018 (2009).

13 Karouzakis, E. et al. Analysis of early changes in DNA methylation in synovial fibroblasts of RA patients before diagnosis. Scientific Reports 8, 7370, doi:10.1038/s41598-018-24240-2 (2018).

14 Nemtsova, M. V. et al. Epigenetic Changes in the Pathogenesis of Rheumatoid Arthritis. Frontiers in Genetics 10, 570, doi:10.3389/fgene.2019.00570 (2019).

15 Nakano, K., Whitaker, J. W., Boyle, D. L., Wang, W. & Firestein, G. S. DNA methylome signature in rheumatoid arthritis. Ann Rheum Dis 72, 110–117, doi:10.1136/annrheumdis-2012-201526 (2013).

16 Doonan, J. et al. The parasitic worm product ES-62 normalises the gut microbiota bone marrow axis in inflammatory arthritis. Nature Communications 10, 1554, doi:10.1038/s41467-019-09361-0 (2019).

17 Pineda, M. A. et al. The parasitic helminth product ES-62 suppresses pathogenesis in collagen-induced arthritis by targeting the interleukin-17-producing cellular network at multiple sites. Arthritis and Rheumatism 64, 3168–3178, doi:10.1002/art.34581 (2012).

18 Pineda, M. A., Rodgers, D. T., Al-Riyami, L., Harnett, W. & Harnett, M. M. ES-62 protects against collagen-induced arthritis by resetting interleukin-22 toward resolution of inflammation in the joints. Arthritis Rheumatol 66, 1492–1503, doi:10.1002/art.38392 (2014).

19 Rodgers, D. T. et al. Protection against collagen-induced arthritis in mice afforded by the parasitic worm product, ES-62, is associated with restoration of the levels of interleukin-10-producing B cells and reduced plasma cell infiltration of the joints. Immunology 141, 457–466, doi:10.1111/imm.12208 (2014).

20 Doonan, J. et al. Protection against arthritis by the parasitic worm project ES-62, and its drug-like small molecule analogues, is associated with inhibition of osteoclastogenesis Frontiers in immunology 9, 1016, doi:10.3389/fimmu.2018.011016 (2018).

21 Fearon, U., Canavan, M., Biniecka, M. & Veale, D. J. Hypoxia, mitochondrial dysfunction and synovial invasiveness in rheumatoid arthritis. Nat Rev Rheumatol 12, 385–397, doi:10.1038/nrrheum.2016.69 (2016).

22 Sharma, J. & Larkin, J., 3rd. Therapeutic Implication of SOCS1 Modulation in the Treatment of Autoimmunity and Cancer. Frontiers in Pharmacology 10, 324, doi:10.3389/fphar.2019.00324 (2019).

23 Malemud, C. J. Negative Regulators of JAK/STAT Signaling in Rheumatoid Arthritis and Osteoarthritis. International Journal of Molecular Sciences 18, doi:10.3390/ijms18030484 (2017).

24 Liang, Y., Xu, W. D., Peng, H., Pan, H. F. & Ye, D. Q. SOCS signaling in autoimmune diseases: molecular mechanisms and therapeutic implications. European Journal of Immunology 44, 1265–1275, doi:10.1002/eji.201344369 (2014).

25 Jiang, M. et al. Dysregulation of SOCS-Mediated Negative Feedback of Cytokine Signaling in Carcinogenesis and Its Significance in Cancer Treatment. Front Immunol 8, 70, doi:10.3389/fimmu.2017.00070 (2017).

26 Zhang, Y. et al. Synergistic effects of interleukin-1beta and interleukin-17A antibodies on collagen-induced arthritis mouse model. Int Immunopharmacol 15, 199–205, doi:10.1016/j.intimp.2012.12.010 (2013).

27 Oike, T. et al. Stat3 as a potential therapeutic target for rheumatoid arthritis. Scientific Reports 7, 10965, doi:10.1038/s41598-017-11233-w (2017).

28 Paunovic, V. & Harnett, M. M. Mitogen-activated protein kinases as therapeutic targets for rheumatoid arthritis. Drugs 73, 101–115, doi:10.1007/s40265-013-0014-6 (2013).

29 Thalhamer, T., McGrath, M. A. & Harnett, M. M. MAPKs and their relevance to arthritis and inflammation. Rheumatology (Oxford) 47, 409–414 (2008).

30 Hirahara, K., Schwartz, D., Gadina, M., Kanno, Y. & O’Shea, J. J. Targeting cytokine signaling in autoimmunity: back to the future and beyond. Current Opinion in Immunology 43, 89–97, doi:10.1016/j.coi.2016.10.001 (2016).

31 Gadina, M., Gazaniga, N., Vian, L. & Furumoto, Y. Small molecules to the rescue: Inhibition of cytokine signaling in immune-mediated diseases. Journal of Autoimmunity, doi:10.1016/j.jaut.2017.06.006 (2017).

32 Mori, T. et al. IL-1beta and TNFalpha-initiated IL-6-STAT3 pathway is critical in mediating inflammatory cytokines and RANKL expression in inflammatory arthritis. International Immunology 23, 701–712, doi:10.1093/intimm/dxr077 (2011).

33 Ganesan, R. & Rasool, M. Interleukin 17 regulates SHP-2 and IL-17RA/STAT-3 dependent Cyr61, IL-23 and GM-CSF expression and RANKL mediated osteoclastogenesis by fibroblast-like synoviocytes in rheumatoid arthritis. Molecular Immunology 91, 134–144, doi:10.1016/j.molimm.2017.09.003 (2017).

34 Morris, K. V., Chan, S. W., Jacobsen, S. E. & Looney, D. J. Small interfering RNA-induced transcriptional gene silencing in human cells. Science (New York, N.Y 305, 1289–1292, doi:10.1126/science.1101372 (2004).

35 Evangelatos, G., Fragoulis, G. E., Koulouri, V. & Lambrou, G. I. MicroRNAs in rheumatoid arthritis: From pathogenesis to clinical impact. Autoimmunity Reviews 18, 102391, doi:10.1016/j.autrev.2019.102391 (2019).

36 Hong, W., Zhang, P., Wang, X., Tu, J. & Wei, W. The Effects of MicroRNAs on Key Signalling Pathways and Epigenetic Modification in Fibroblast-Like Synoviocytes of Rheumatoid Arthritis. Mediators Inflamm 2018, 9013124, doi:10.1155/2018/9013124 (2018).

37 Wu, H., Chen, Y., Zhu, H., Zhao, M. & Lu, Q. The Pathogenic Role of Dysregulated Epigenetic Modifications in Autoimmune Diseases. Frontiers in Immunology 10, 2305, doi:10.3389/fimmu.2019.02305 (2019).

38 Sharma, A. R., Sharma, G., Lee, S. S. & Chakraborty, C. miRNA-Regulated Key Components of Cytokine Signaling Pathways and Inflammation in Rheumatoid Arthritis. Med Res Rev 36, 425–439, doi:10.1002/med.21384 (2016).

39 Su, L. C., Huang, A. F., Jia, H., Liu, Y. & Xu, W. D. Role of microRNA-155 in rheumatoid arthritis. International Journal of Rheumatic Diseases 20, 1631–1637, doi:10.1111/1756-185X.13202 (2017).

40 Alivernini, S. et al. MicroRNA-155-at the Critical Interface of Innate and Adaptive Immunity in Arthritis. Frontiers in Immunology 8, 1932, doi:10.3389/fimmu.2017.01932 (2017).

41 Stanczyk, J. et al. Altered expression of MicroRNA in synovial fibroblasts and synovial tissue in rheumatoid arthritis. Arthritis and Rheumatism 58, 1001–1009, doi:10.1002/art.23386 (2008).

42 Abdollahi-Roodsaz, S., van de Loo, F. A. & van den Berg, W. B. Trapped in a vicious loop: Toll-like receptors sustain the spontaneous cytokine production by rheumatoid synovium. Arthritis Research & Therapy 13, 105, doi:ar3287 [pii] 10.1186/ar3287 (2011).

43 Arleevskaya, M. I., Larionova, R. V., Brooks, W. H., Bettacchioli, E. & Renaudineau, Y. Toll-Like Receptors, Infections, and Rheumatoid Arthritis. Clinical Reviews in Allergy & Immunology 58, 172–181, doi:10.1007/s12016-019-08742-z (2020).

44 Bluml, S. et al. Essential role of microRNA-155 in the pathogenesis of autoimmune arthritis in mice. Arthritis and Rheumatism 63, 1281–1288, doi:10.1002/art.30281 (2011).

45 Kurowska-Stolarska, M. et al. MicroRNA-155 as a proinflammatory regulator in clinical and experimental arthritis. Proceedings of the National Academy of Sciences of the United States of America 108, 11193–11198, doi:10.1073/pnas.1019536108 (2011).

46 Stanczyk, J. et al. Altered expression of microRNA-203 in rheumatoid arthritis synovial fibroblasts and its role in fibroblast activation. Arthritis and Rheumatism 63, 373–381, doi:10.1002/art.30115 (2011).

47 Long, L. et al. Upregulated microRNA-155 expression in peripheral blood mononuclear cells and fibroblast-like synoviocytes in rheumatoid arthritis. Clinical & Developmental Immunology 2013, 296139, doi:10.1155/2013/296139 (2013).

48 Liu, N., Feng, X., Wang, W., Zhao, X. & Li, X. Paeonol protects against TNF- alpha-induced proliferation and cytokine release of rheumatoid arthritis fibroblast-like synoviocytes by upregulating FOXO3 through inhibition of miR- 155 expression. Inflamm Res 66, 603–610, doi:10.1007/s00011-017-1041-7 (2017).

49 Xie, Z., et al. PU.1 attenuates TNFalphainduced proliferation and cytokine release of rheumatoid arthritis fibroblastlike synoviocytes by regulating miR155 activity. Molecular Medicine Reports 17, 8349–8356, doi:10.3892/mmr.2018.8920 (2018).

50 Wang, Y. et al. miR-155 promotes fibroblast-like synoviocyte proliferation and inflammatory cytokine secretion in rheumatoid arthritis by targeting FOXO3a. Experimental and Therapeutic Medicine 19, 1288–1296, doi:10.3892/etm.2019.8330 (2020).

51 Brandstetter, B. et al. FOXO3 is involved in the tumor necrosis factor-driven inflammatory response in fibroblast-like synoviocytes. Lab Invest 99, 648–658, doi:10.1038/s41374-018-0184-7 (2019).

52 Nakagawa, R. et al. MicroRNA-155 controls affinity-based selection by protecting c-MYC+ B cells from apoptosis. The Journal of Clinical Investigation 126, 377–388, doi:10.1172/JCI82914 (2016).

53 Thai, T. H. et al. Regulation of the germinal center response by microRNA-155. Science (New York, N.Y 316, 604–608 (2007).

54 Li, X., Tian, F. & Wang, F. Rheumatoid arthritis-associated microRNA-155 targets SOCS1 and upregulates TNF-alpha and IL-1beta in PBMCs. International Journal of Molecular Sciences 14, 23910–23921, doi:10.3390/ijms141223910 (2013).

55 Gujar, H., Weisenberger, D. J. & Liang, G. The Roles of Human DNA Methyltransferases and Their Isoforms in Shaping the Epigenome. Genes 10, doi:10.3390/genes10020172 (2019).

56 Brenet, F. et al. DNA methylation of the first exon is tightly linked to transcriptional silencing. PLoS ONE 6, e14524, doi:10.1371/journal.pone.0014524 (2011).

57 Jeltsch, A. & Jurkowska, R. Z. New concepts in DNA methylation. Trends in Biochemical Sciences 39, 310–318, doi:10.1016/j.tibs.2014.05.002 (2014).

58 Lev Maor, G., Yearim, A. & Ast, G. The alternative role of DNA methylation in splicing regulation. Trends Genet 31, 274–280, doi:10.1016/j.tig.2015.03.002 (2015).

59 Tirado-Magallanes, R., Rebbani, K., Lim, R., Pradhan, S. & Benoukraf, T. Whole genome DNA methylation: beyond genes silencing. Oncotarget 8, 5629–5637, doi:10.18632/oncotarget.13562 (2017).

60 Janssens, S., Burns, K., Tschopp, J. & Beyaert, R. Regulation of interleukin-1- and lipopolysaccharide-induced NF-kappaB activation by alternative splicing of MyD88. Curr Biol 12, 467–471 (2002).

61 Janssens, S., Burns, K., Vercammen, E., Tschopp, J. & Beyaert, R. MyD88S, a splice variant of MyD88, differentially modulates NF-κB- and AP-1- dependent gene expression. FEBS Letters 548, 103–107, doi:10.1016/s0014-5793(03)00747-6 (2003).

62 Biamonti, G., Infantino, L., Gaglio, D. & Amato, A. An Intricate Connection between Alternative Splicing and Phenotypic Plasticity in Development and Cancer. Cells 9, doi:10.3390/cells9010034 (2019).

63 Collins, I. & Wann, A. K. T. Regulation of the Extracellular Matrix by Ciliary Machinery. Cells 9, doi:10.3390/cells9020278 (2020).

64 Kopinke, D., Norris, A. M. & Mukhopadhyay, S. Developmental and regenerative paradigms of cilia regulated hedgehog signaling. Semin Cell Dev Biol, doi:10.1016/j.semcdb.2020.05.029 (2020).

65 Boukhalfa, A., Miceli, C., Avalos, Y., Morel, E. & Dupont, N. Interplay between primary cilia, ubiquitin-proteasome system and autophagy. Biochimie 166, 286–292, doi:10.1016/j.biochi.2019.06.009 (2019).

66 Martin-Hurtado, A., Lastres-Becker, I., Cuadrado, A. & Garcia-Gonzalo, F. R. NRF2 and Primary Cilia: An Emerging Partnership. Antioxidants (Basel) 9, doi:10.3390/antiox9060475 (2020).

67 Tao, F., Jiang, T., Tao, H., Cao, H. & Xiang, W. Primary cilia: Versatile regulator in cartilage development. Cell Prolif 53, e12765, doi:10.1111/cpr.12765 (2020).

68 Teves, M. E., Strauss, J. F., 3rd, Sapao, P., Shi, B. & Varga, J. The Primary Cilium: Emerging Role as a Key Player in Fibrosis. Curr Rheumatol Rep 21, 29, doi:10.1007/s11926-019-0822-0 (2019).

69 Eason, R. J., et al. The helminth product, ES-62 modulates dendritic cell responses by inducing the selective autophagolysosomal degradation of TLR- transducers, as exemplified by PKCdelta. Scientific Reports 6, 37276, doi:10.1038/srep37276 (2016).

70 Lee, H. M. et al. Autophagy negatively regulates keratinocyte inflammatory responses via scaffolding protein p62/SQSTM1. J Immunol 186, 1248–1258, doi:10.4049/jimmunol.1001954 (2011).

71 Klionsky, D. J. et al. Guidelines for the use and interpretation of assays for monitoring autophagy (3rd edition). Autophagy 12, 1–222, doi:10.1080/15548627.2015.1100356 (2016).

72 O’Rourke, J. G. et al. C9orf72 is required for proper macrophage and microglial function in mice. *Science (New York*, N.Y 351, 1324–1329, doi:10.1126/science.aaf1064 (2016).

73 Deretic, V., Saitoh, T. & Akira, S. Autophagy in infection, inflammation and immunity. Nature Reviews Immunology 13, 722–737, doi:10.1038/nri3532 (2013).

74 Rzepecka, J. et al. Prophylactic and therapeutic treatment with a synthetic analogue of a parasitic worm product prevents experimental arthritis and inhibits IL-1beta production via NRF2-mediated counter-regulation of the inflammasome. Journal of Autoimmunity 60, 59–73, doi:10.1016/j.jaut.2015.04.005 (2015).

75 Roeleveld, D. M. & Koenders, M. I. The role of the Th17 cytokines IL-17 and IL-22 in Rheumatoid Arthritis pathogenesis and developments in cytokine immunotherapy. Cytokine 74, 101–107, doi:10.1016/j.cyto.2014.10.006 (2015).

76 Nakae, S., Nambu, A., Sudo, K. & Iwakura, Y. Suppression of immune induction of collagen-induced arthritis in IL-17-deficient mice. Journal of Immunology 171, 6173–6177 (2003).

77 Pineda, M. A., Lumb, F., Harnett, M. M. & Harnett, W. ES-62, a therapeutic anti-inflammatory agent evolved by the filarial nematode Acanthocheilonema viteae. Molecular and Biochemical Parasitology 194, 1–8, doi:10.1016/j.molbiopara.2014.03.003 (2014).

78 Alexander, W. S. et al. SOCS1 is a critical inhibitor of interferon gamma signaling and prevents the potentially fatal neonatal actions of this cytokine. Cell 98, 597–608 (1999).

79 Dolhain, R. J. et al. Increased expression of interferon (IFN)-gamma together with IFN-gamma receptor in the rheumatoid synovial membrane compared with synovium of patients with osteoarthritis. Br J Rheumatol 35, 24–32 (1996).

80 T Karonitsch, K. D., R Byrne, B Niedereiter, E Cetin, A Wanivenhaus, C Scheinecker, J S Smolen, H P Kiener. IFN-gamma promotes fibroblast-like synoviocytes motility. Ann Rheum Dis 69 (2010).

81 Marshall, F. A., Grierson, A. M., Garside, P., Harnett, W. & Harnett, M. M. ES-62, an immunomodulator secreted by filarial nematodes, suppresses clonal expansion and modifies effector function of heterologous antigen-specific T cells in vivo. J Immunol 175, 5817–5826 (2005).

82 Goodridge, H. S. et al. Modulation of macrophage cytokine production by ES-62, a secreted product of the filarial nematode Acanthocheilonema viteae. J Immunol 167, 940–945 (2001).

83 Fang, S. et al. MiR-455 targeting SOCS3 improve liver lipid disorders in diabetic mice. Adipocyte 9, 179–188, doi:10.1080/21623945.2020.1749495 (2020).

84 Dey, A., Varelas, X. & Guan, K. L. Targeting the Hippo pathway in cancer, fibrosis, wound healing and regenerative medicine. Nat Rev Drug Discov, doi:10.1038/s41573-020-0070-z (2020).

85 Yoshida, G. J. Regulation of heterogeneous cancer-associated fibroblasts: the molecular pathology of activated signaling pathways. J Exp Clin Cancer Res 39, 112, doi:10.1186/s13046-020-01611-0 (2020).

86 Harnett, M. M., Melendez, A. J. & Harnett, W. The therapeutic potential of the filarial nematode-derived immunodulator, ES-62 in inflammatory disease. Clinical and Experimental Immunology 159, 256–267, doi:CEI4064 [pii] 10.1111/j.1365-2249.2009.04064.x (2010).

87 Pan, J. A. et al. miR-146a attenuates apoptosis and modulates autophagy by targeting TAF9b/P53 pathway in doxorubicin-induced cardiotoxicity. Cell Death & Disease 10, 668, doi:10.1038/s41419-019-1901-x (2019).

88 Zhang, Q. et al. The MAP3K13-TRIM25-FBXW7alpha axis affects c-Myc protein stability and tumor development. Cell Death and Differentiation 27, 420–433, doi:10.1038/s41418-019-0363-0 (2020).

89 Zhang, L., Li, H., Yuan, M., Li, M. & Zhang, S. Cervical Cancer Cells-Secreted Exosomal microRNA-221-3p Promotes Invasion, Migration and Angiogenesis of Microvascular Endothelial Cells in Cervical Cancer by Down-Regulating MAPK10 Expression. Cancer Manag Res 11, 10307–10319, doi:10.2147/CMAR.S221527 (2019).

90 Li, H., Zuo, J. & Tang, W. Phosphodiesterase-4 Inhibitors for the Treatment of Inflammatory Diseases. Frontiers in Pharmacology 9, 1048, doi:10.3389/fphar.2018.01048 (2018).

91 Tschaikner, P., Enzler, F., Torres-Quesada, O., Aanstad, P. & Stefan, E. Hedgehog and Gpr161: Regulating cAMP Signaling in the Primary Cilium. Cells 9, doi:10.3390/cells9010118 (2020).

92 Zhu, S. et al. Sonic Hedgehog Regulates Proliferation, Migration and Invasion of Synoviocytes in Rheumatoid Arthritis via JNK Signaling. Frontiers in Immunology 11, 1300, doi:10.3389/fimmu.2020.01300 (2020).

93 Comertpay, B. & Gov, E. Identification of key biomolecules in rheumatoid arthritis through the reconstruction of comprehensive disease-specific biological networks. Autoimmunity 53, 156–166, doi:10.1080/08916934.2020.1722107 (2020).

94 Rodgers, D. T. et al. The Parasitic Worm Product ES-62 Targets Myeloid Differentiation Factor 88-Dependent Effector Mechanisms to Suppress Antinuclear Antibody Production and Proteinuria in MRL/lpr Mice. Arthritis Rheumatol 67, 1023–1035, doi:10.1002/art.39004 (2015).

95 Goodridge, H. S. et al. In vivo exposure of murine dendritic cell and macrophage bone marrow progenitors to the phosphorylcholine-containing filarial nematode glycoprotein ES-62 polarizes their differentiation to an anti-inflammatory phenotype. Immunology 113, 491–498 (2004).

96 Mei, X. F. et al. DNA methylation and hydroxymethylation profiles reveal possible role of highly methylated TLR signaling on Fasciola gigantica excretory/secretory products (FgESPs) modulation of buffalo dendritic cells. Parasites & Vectors 12, 358, doi:10.1186/s13071-019-3615-4 (2019).

97 Cook, P. C. et al. A dominant role for the methyl-CpG-binding protein Mbd2 in controlling Th2 induction by dendritic cells. Nature Communications 6, 6920, doi:10.1038/ncomms7920 (2015).

98 Jones, G. R. et al. The Methyl-CpG-Binding Protein Mbd2 Regulates Susceptibility to Experimental Colitis via Control of CD11c(+) Cells and Colonic Epithelium. Frontiers in Immunology 11, 183, doi:10.3389/fimmu.2020.00183 (2020).

99 Jin, S. G., Jiang, C. L., Rauch, T., Li, H. & Pfeifer, G. P. MBD3L2 interacts with MBD3 and components of the NuRD complex and can oppose MBD2-MeCP1-mediated methylation silencing. The Journal of Biological Chemistry 280, 12700–12709, doi:10.1074/jbc.M413492200 (2005).

100 Ginder, G. D. & Williams, D. C., Jr. Readers of DNA methylation, the MBD family as potential therapeutic targets. Pharmacology & Therapeutics 184, 98–111, doi:10.1016/j.pharmthera.2017.11.002 (2018).

101 Mota, E. A. et al. Epigenetic and parasitological parameters are modulated in EBi3-/- mice infected with Schistosoma mansoni. PLoS Neglected Tropical Diseases 14, e0008080, doi:10.1371/journal.pntd.0008080 (2020).

102 DiNardo, A. R. et al. Schistosomiasis Induces Persistent DNA Methylation and Tuberculosis-Specific Immune Changes. J Immunol, doi:10.4049/jimmunol.1800101 (2018).

103 Klar, K. et al. Chronic schistosomiasis during pregnancy epigenetically reprograms T-cell differentiation in offspring of infected mothers. European Journal of Immunology 47, 841–847, doi:10.1002/eji.201646836 (2017).

104 Dewals, B. G., Layland, L. E., Prazeres da Costa, C. & Horsnell, W. G. Maternal helminth infections and the shaping of offspring immunity. Parasite Immunology 41, e12599, doi:10.1111/pim.12599 (2019).

105 McInnes, I. B. et al. A Novel Therapeutic Approach Targeting Articular Inflammation Using the Filarial Nematode-Derived Phosphorylcholine-Containing Glycoprotein ES-62. The Journal of Immunology 171, 2127–2133, doi:10.4049/jimmunol.171.4.2127 (2003).

106 Maria Armaka, V. G., Dimitris Kontoyiannis & George Kollias. A standardized protocol for the isolation and culture of normal and arthritogenic murine synovial fibroblasts. Protocol Exchange 102 (2009).

107 Gelse, K. et al. Role of hypoxia-inducible factor 1 alpha in the integrity of articular cartilage in murine knee joints. Arthritis Research & Therapy 10, R111, doi:10.1186/ar2508 (2008).

108 Radu, M. & Chernoff, J. An in vivo assay to test blood vessel permeability. J Vis Exp, e50062, doi:10.3791/50062 (2013).

109 Wilson, E. H. et al. Hyporesponsiveness of murine B lymphocytes exposed to the filarial nematode secreted product ES-62 in vivo. Immunol 109, 238–245 (2003).

110 Lumb, F. E. et al. Dendritic cells provide a therapeutic target for synthetic small molecule analogues of the parasitic worm product, ES-62. Scientific Reports 7, 1704, doi:10.1038/s41598-017-01651-1 (2017).

111 Marzi, M. J. a. N., F. Flexible, efficient miRNA detection using the miScript PCR System. Qiagen.

112 Kurowska-Stolarska, M. e. a. MicroRNA-155 as a proinflammatory regulator in clinical and experimental arthritis. Proceedings of the National Academy of Sciences of the United States of America 108, 11193–11198 (2011).

113 Nakano, K., Boyle, D. L. & Firestein, G. S. Regulation of DNA methylation in rheumatoid arthritis synoviocytes. J Immunol 190, 1297–1303, doi:10.4049/jimmunol.1202572 (2013).

## Cited References

1 Ormseth, M. J. et al. Development and Validation of a MicroRNA Panel to Differentiate Between Patients with Rheumatoid Arthritis or Systemic Lupus Erythematosus and Controls. The Journal of Rheumatology 47, 188–196, doi:10.3899/jrheum.181029 (2020).

2 Evangelatos, G., Fragoulis, G. E., Koulouri, V. & Lambrou, G. I. MicroRNAs in rheumatoid arthritis: From pathogenesis to clinical impact. Autoimmunity Reviews 18, 102391, doi:10.1016/j.autrev.2019.102391 (2019).

3 Hong, W., Zhang, P., Wang, X., Tu, J. & Wei, W. The Effects of MicroRNAs on Key Signalling Pathways and Epigenetic Modification in Fibroblast-Like Synoviocytes of Rheumatoid Arthritis. Mediators Inflamm 2018, 9013124, doi:10.1155/2018/9013124 (2018).

4 Sharma, A. R., Sharma, G., Lee, S. S. & Chakraborty, C. miRNA-Regulated Key Components of Cytokine Signaling Pathways and Inflammation in Rheumatoid Arthritis. Med Res Rev 36, 425–439, doi:10.1002/med.21384 (2016).

5 Su, L. C., Huang, A. F., Jia, H., Liu, Y. & Xu, W. D. Role of microRNA-155 in rheumatoid arthritis. International Journal of Rheumatic Diseases 20, 1631–1637, doi:10.1111/1756-185X.13202 (2017).

6 Park, S. H. et al. BATF regulates collagen-induced arthritis by regulating T helper cell differentiation. Arthritis Research & Therapy 20, 161, doi:10.1186/s13075-018-1658-0 (2018).

7 Webster, A. P. et al. Increased DNA methylation variability in rheumatoid arthritis-discordant monozygotic twins. Genome Med 10, 64, doi:10.1186/s13073-018-0575-9 (2018).

8 Plestilova, L. et al. Expression and Regulation of PIWIL-Proteins and PIWI-Interacting RNAs in Rheumatoid Arthritis. PLoS ONE 11, e0166920, doi:10.1371/journal.pone.0166920 (2016).

9 Chang, X. et al. CD38 and E2F transcription factor 2 have uniquely increased expression in rheumatoid arthritis synovial tissues. Clinical and Experimental Immunology 176, 222–231, doi:10.1111/cei.12268 (2014).

10 Janczi, T. et al. ADAM15 in Apoptosis Resistance of Synovial Fibroblasts: Converting Fas/CD95 Death Signals Into the Activation of Prosurvival Pathways by Calmodulin Recruitment. Arthritis Rheumatol 71, 63–72, doi:10.1002/art.40667 (2019).

11 Devauchelle, V. et al. DNA microarray allows molecular profiling of rheumatoid arthritis and identification of pathophysiological targets. Genes and Immunity 5, 597–608, doi:10.1038/sj.gene.6364132 (2004).

12 Lee, E. J. et al. Interleukin-33 acts as a transcriptional repressor and extracellular cytokine in fibroblast-like synoviocytes in patients with rheumatoid arthritis. Cytokine 77, 35–43, doi:10.1016/j.cyto.2015.10.005 (2016).

13 Shi, L. J. et al. Elevated Levels of Soluble ST2 were Associated with Rheumatoid Arthritis Disease Activity and Ameliorated Inflammation in Synovial Fibroblasts. Chin Med J (Engl) 131, 316–322, doi:10.4103/0366-6999.223847 (2018).

14 Zhang, Y. et al. LAIR-1 shedding from human fibroblast-like synoviocytes in rheumatoid arthritis following TNF-alpha stimulation. Clinical and Experimental Immunology 192, 193–205, doi:10.1111/cei.13100 (2018).

15 Fu, D. et al. Role of p21-activated kinase 1 in regulating the migration and invasion of fibroblast-like synoviocytes from rheumatoid arthritis patients. Rheumatology (Oxford) 51, 1170–1180, doi:10.1093/rheumatology/kes031 (2012).

16 Chen, W. et al. Apremilast Ameliorates Experimental Arthritis via Suppression of Th1 and Th17 Cells and Enhancement of CD4(+)Foxp3(+) Regulatory T Cells Differentiation. Frontiers in Immunology 9, 1662, doi:10.3389/fimmu.2018.01662 (2018).

17 Kobayashi, K., Suda, T., Manabe, H. & Miki, I. Administration of PDE4 inhibitors suppressed the pannus-like inflammation by inhibition of cytokine production by macrophages and synovial fibroblast proliferation. Mediators Inflamm 2007, 58901, doi:10.1155/2007/58901 (2007).

18 McCann, F. E. et al. Apremilast, a novel PDE4 inhibitor, inhibits spontaneous production of tumour necrosis factor-alpha from human rheumatoid synovial cells and ameliorates experimental arthritis. Arthritis Research & Therapy 12, R107, doi:10.1186/ar3041 (2010).

19 Yang, B. et al. miR-124a inhibits the proliferation and inflammation in rheumatoid arthritis fibroblast-like synoviocytes via targeting PIK3/NF-kappaB pathway. Cell Biochem Funct 37, 208–215, doi:10.1002/cbf.3386 (2019).

20 Miao, C. et al. CUL4B promotes the pathology of adjuvant-induced arthritis in rats through the canonical Wnt signaling. J Mol Med (Berl) 96, 495–511, doi:10.1007/s00109-018-1635-8 (2018).

21 He, S. D. et al. Oridonin suppresses autophagy and survival in rheumatoid arthritis fibroblast-like synoviocytes. Pharm Biol 58, 146–151, doi:10.1080/13880209.2020.1711783 (2020).

22 Dinesh, P. & Rasool, M. Berberine mitigates IL-21/IL-21R mediated autophagic influx in fibroblast-like synoviocytes and regulates Th17/Treg imbalance in rheumatoid arthritis. Apoptosis 24, 644–661, doi:10.1007/s10495-019-01548-6 (2019).

23 Deng, H. et al. Effects of daphnetin on the autophagy signaling pathway of fibroblast-like synoviocytes in rats with collagen-induced arthritis (CIA) induced by TNF-alpha. Cytokine 127, 154952, doi:10.1016/j.cyto.2019.154952 (2020).

24 Sugawara, E. et al. Autophagy promotes citrullination of VIM (vimentin) and its interaction with major histocompatibility complex class II in synovial fibroblasts. Autophagy 16, 946–955, doi:10.1080/15548627.2019.1664144 (2020).

25 Ikegawa, S. Expression, regulation and function of asporin, a susceptibility gene in common bone and joint diseases. Current Medicinal Chemistry 15, 724–728, doi:10.2174/092986708783885237 (2008).

26 Torres, B. et al. Asporin repeat polymorphism in rheumatoid arthritis. Ann Rheum Dis 66, 118–120, doi:10.1136/ard.2006.055426 (2007).

27 Volin, M. V., Shahrara, S., Haines, G. K., 3rd, Woods, J. M. & Koch, A. E. Expression of mucin 3 and mucin 5AC in arthritic synovial tissue. Arthritis and Rheumatism 58, 46–52, doi:10.1002/art.23174 (2008).

28 Anaparti, V., Agarwal, P., Smolik, I., Mookherjee, N. & Elgabalawy, H. Whole Blood Targeted Bisulfite Sequencing Validates Differential Methylation in C6ORF10 gene of Patients with Rheumatoid Arthritis. The Journal of Rheumatology, doi:10.3899/jrheum.190376 (2019).

29 Yokoyama, W. et al. Abrogation of CC chemokine receptor 9 ameliorates collagen-induced arthritis of mice. Arthritis Research & Therapy 16, 445, doi:10.1186/s13075-014-0445-9 (2014).

30 Chen, Z. et al. Characterising the expression and function of CCL28 and its corresponding receptor, CCR10, in RA pathogenesis. Ann Rheum Dis 74, 1898–1906, doi:10.1136/annrheumdis-2013-204530 (2015).

31 Redelinghuys, P. et al. MICL controls inflammation in rheumatoid arthritis. Ann Rheum Dis 75, 1386–1391, doi:10.1136/annrheumdis-2014-206644 (2016).

32 Sawai, H., Park, Y. W., He, X., Goronzy, J. J. & Weyand, C. M. Fractalkine mediates T cell-dependent proliferation of synovial fibroblasts in rheumatoid arthritis. Arthritis and Rheumatism 56, 3215–3225, doi:10.1002/art.22919 (2007).

33 Laragione, T., Brenner, M., Sherry, B. & Gulko, P. S. CXCL10 and its receptor CXCR3 regulate synovial fibroblast invasion in rheumatoid arthritis. Arthritis and Rheumatism 63, 3274–3283, doi:10.1002/art.30573 (2011).

34 Li, C. H. et al. CXCL16 upregulates RANKL expression in rheumatoid arthritis synovial fibroblasts through the JAK2/STAT3 and p38/MAPK signaling pathway. Inflamm Res 65, 193–202, doi:10.1007/s00011-015-0905-y (2016).

35 Marotte, H. et al. Blocking of interferon regulatory factor 1 reduces tumor necrosis factor alpha-induced interleukin-18 bioactivity in rheumatoid arthritis synovial fibroblasts by induction of interleukin-18 binding protein a: role of the nuclear interferon regulatory factor 1-NF-kappaB-c-jun complex. Arthritis and Rheumatism 63, 3253–3262, doi:10.1002/art.30583 (2011).

36 Chen, J., Caspi, R. R. & Chong, W. P. IL-20 receptor cytokines in autoimmune diseases. Journal of Leukocyte Biology 104, 953–959, doi:10.1002/JLB.MR1117-471R (2018).

37 Gong, F. et al. Interleukin-27 as a potential therapeutic target for rheumatoid arthritis: has the time come? Clin Rheumatol 32, 1425–1428, doi:10.1007/s10067-013-2341-0 (2013).

38 Ye, S. et al. Variation in the matrix metalloproteinase-3, -7, -12 and -13 genes is associated with functional status in rheumatoid arthritis. Int J Immunogenet 34, 81–85, doi:10.1111/j.1744-313X.2007.00664.x (2007).

39 Knevel, R. et al. Identification of a genetic variant for joint damage progression in autoantibody-positive rheumatoid arthritis. Ann Rheum Dis 73, 2038–2046, doi:10.1136/annrheumdis-2013-204050 (2014).

40 Elshabrawy, H. A., Essani, A. E., Szekanecz, Z., Fox, D. A. & Shahrara, S. TLRs, future potential therapeutic targets for RA. Autoimmunity Reviews 16, 103–113, doi:10.1016/j.autrev.2016.12.003 (2017).

41 Kim, K. W. et al. Toll-like receptor 7 regulates osteoclastogenesis in rheumatoid arthritis. J Biochem 166, 259–270, doi:10.1093/jb/mvz033 (2019).

42 Liu, R. et al. Adiponectin promotes fibroblast-like synoviocytes producing IL-6 to enhance T follicular helper cells response in rheumatoid arthritis. Clinical and Experimental Rheumatology 38, 11–18 (2020).

43 Zhao, Y. et al. PGK1, a glucose metabolism enzyme, may play an important role in rheumatoid arthritis. Inflamm Res 65, 815–825, doi:10.1007/s00011-016-0965-7 (2016).

44 Huang, G. et al. Loss of core fucosylation in both ST6GAL1 and its substrate enhances glycoprotein sialylation in mice. The Biochemical Journal 477, 1179–1201, doi:10.1042/BCJ20190789 (2020).

45 Lauc, G. et al. Loci associated with N-glycosylation of human immunoglobulin G show pleiotropy with autoimmune diseases and haematological cancers. PLoS Genet 9, e1003225, doi:10.1371/journal.pgen.1003225 (2013).

46 Gonzalez-Rey, E., Chorny, A., Del Moral, R. G., Varela, N. & Delgado, M. Therapeutic effect of cortistatin on experimental arthritis by downregulating inflammatory and Th1 responses. Ann Rheum Dis 66, 582–588, doi:10.1136/ard.2006.062703 (2007).

